# Phase-Based Frequency Tracking and Analysis of Vocalisations in the Genus Saxicola

**DOI:** 10.64898/2026.10.14.759137

**Authors:** F. P. L. Strijbos

## Abstract

Acoustic analysis provides valuable tools for studying bird vocalisations, but quantitative characterisation of call structure remains challenging when frequency and amplitude vary jointly over time. We present a phase-based frequency tracking method that represents amplitude as a function of instantaneous frequency and use it to characterise alarm calls in the genus *Saxicola*. The method enables the extraction of acoustic features describing the frequency trajectory and temporal structure of individual calls. We apply the method to a dataset of 801 calls from 49 recordings representing 15 species, and use principal component analysis (PCA) and robust PCA to investigate patterns of acoustic variation across species, recordings and geographic regions. The results demonstrate the feasibility of the approach and reveal structured patterns of acoustic similarity and variation among the calls analysed. Robust PCA concentrates the variance of the low-rank representation into fewer principal components than conventional PCA applied to the original feature matrix, providing a complementary representation of the dominant acoustic structure. The observed patterns illustrate the potential of quantitative acoustic characterisation for comparative studies of vocal variation. However, uneven species representation and uncertainty in individual identity and taxonomic labels limit the biological conclusions that can be drawn from the present dataset. The method provides a foundation for further acoustic research, in which broader sampling and integration with independent genetic and morphological evidence could help investigate variation within and among species.

## 1 INTRODUCTION

Accurate species identification is crucial for biodiversity research and ecosystem monitoring and conservation (Sethi et al., 2020). For many species, especially those that are difficult to observe directly, acoustic recordings offer an efficient means of detecting and identifying individuals in their natural habitat. Advances in automated acoustic monitoring have further expanded the potential to survey species over vast spatial and temporal scales. However, reliable identification hinges on extracting meaningful characteristics from complex acoustic signals. Bird vocalisations are particularly well suited to acoustic analysis. Many calls and songs feature distinctive patterns of frequency, amplitude and temporal structure. These patterns provide valuable information not only for species identification, but also for exploring geographic and intraspecific variation in vocal traits. Consequently, the representation and quantification of acoustic signals can significantly influence the biological information that can be extracted from them. In ornithology, spectrograms are commonly analysed for this purpose. A spectrogram represents a vocalisation in three dimensions—time, frequency and amplitude—but conventional contour extraction largely reduces this information to a two-dimensional time–frequency representation. Amplitude relative to a threshold primarily determines whether a time– frequency bin is retained in the extracted contour (Buck and Tyack, 1993; Mellinger et al., 2011). In practice, however, call amplitude may vary substantially, from a local maximum to values well below such a threshold. Moreover, this approach emphasizes the relationship between time and frequency rather than the relationship between amplitude and frequency. The latter may provide an additional avenue for investigating the physical processes underlying sound production. In this paper, we present a new method for analysing amplitude as a function of frequency, where frequency itself varies with time. We compare this approach with conventional analysis of frequency as a function of time, in which amplitude plays primarily a qualitative role.

If the amplitude–frequency relationship contains biologically meaningful information, then a method that explicitly preserves this relationship should allow acoustic features to be extracted that are not captured by a conventional frequency contour.

## 2 METHODOLOGY

### 2.1 Phase-based Frequency Tracking

The proposed method uses a reference frequency, or ‘anchor frequency’, to extract the instantaneous frequency and amplitude of a vocalisation. The anchor frequency can be selected as a characteristic frequency for a species, estimated from the recorded signal, for example by counting zero crossings, or obtained from a previously determined time–frequency contour.

The recorded signal is mixed with a sinusoid at the anchor frequency. This shifts the signal’s frequency content relative to the anchor, translating the frequency of interest towards base-band and producing a difference frequency that can be readily analysed. A second mixing operation, using a reference sinusoid shifted by a quarter of a cycle, produces a corresponding signal in quadrature. Together, these two signals provide the information needed to determine the instantaneous phase of the difference-frequency signal.

The rate at which this phase changes gives the instantaneous frequency relative to the anchor. By adding the anchor frequency, the instantaneous frequency of the original vocalisation can then be recovered. The two quadrature signals also provide an estimate of the instantaneous amplitude. In this way, the method simultaneously tracks the frequency and amplitude of the vocalisation, allowing their relationship to be examined throughout the call.

For a mathematical derivation of the phase-based frequency tracking method, we refer to Appendix A.1. Appendix A.2 includes a comparison with Fourier-based frequency estimation on noise-free and noisy synthetic data. The smoothing inherent to the Short-Time Fourier Transform (STFT) depends on the choice of window duration. This window duration is typically selected to achieve an appropriate time–frequency resolution rather than specifically for noise suppression. In the proposed method, frequency tracking and smoothing are separated: the instantaneous frequency is first obtained from the phase, after which a Kalman filter can be used to estimate its underlying trend. This provides explicit control over the degree of smoothing without changing the frequency-tracking procedure itself.

Unlike a conventional frequency contour, which primarily describes how frequency changes with time, the proposed method simultaneously estimates frequency and amplitude. It therefore retains the amplitude–frequency structure of the vocalisation for subsequent analysis.

### 2.2 Call segmentation and parameterisation of the amplitude-frequency trends

To investigate the amplitude–frequency structure of individual calls, each call was divided into three sections: onset, sustain and decay. The maximum amplitude was first identified. The onset was defined as the portion of the call preceding the maximum amplitude and beginning when the amplitude exceeded a threshold relative to the maximum. The end of the call was determined from the estimated noise floor, calculated from the final part of the recording.

The sustain was defined as the approximately constant-amplitude region surrounding the maximum amplitude. Rather than imposing a fixed lower amplitude boundary, the lower boundary of the sustain was progressively varied. For each candidate sustain region, the frequency–amplitude relationship during the onset and decay was described by a polynomial, while the frequency during the sustain was represented by its mean value. The candidate segmentation giving the smallest overall frequency standard error across the three sections was selected.

This procedure therefore identifies the three sections from the structure of the amplitude–frequency relationship itself. The resulting onset and decay sections describe how frequency changes with amplitude, whereas the sustain represents a region of approximately constant amplitude, with its frequency summarised by the mean value. These relationships were subsequently parameterised to form the feature vector used for the multivariate analysis. The features were selected to capture frequency level, frequency variation, frequency–amplitude geometry, and temporal duration while avoiding redundant descriptors. Each element of this vector is described in Table 1. Check marks indicate retained features; crosses indicate features excluded during feature selection. A more detailed description of the feature vector derivation can be found in Appendix B.2.

**TABLE 1.**
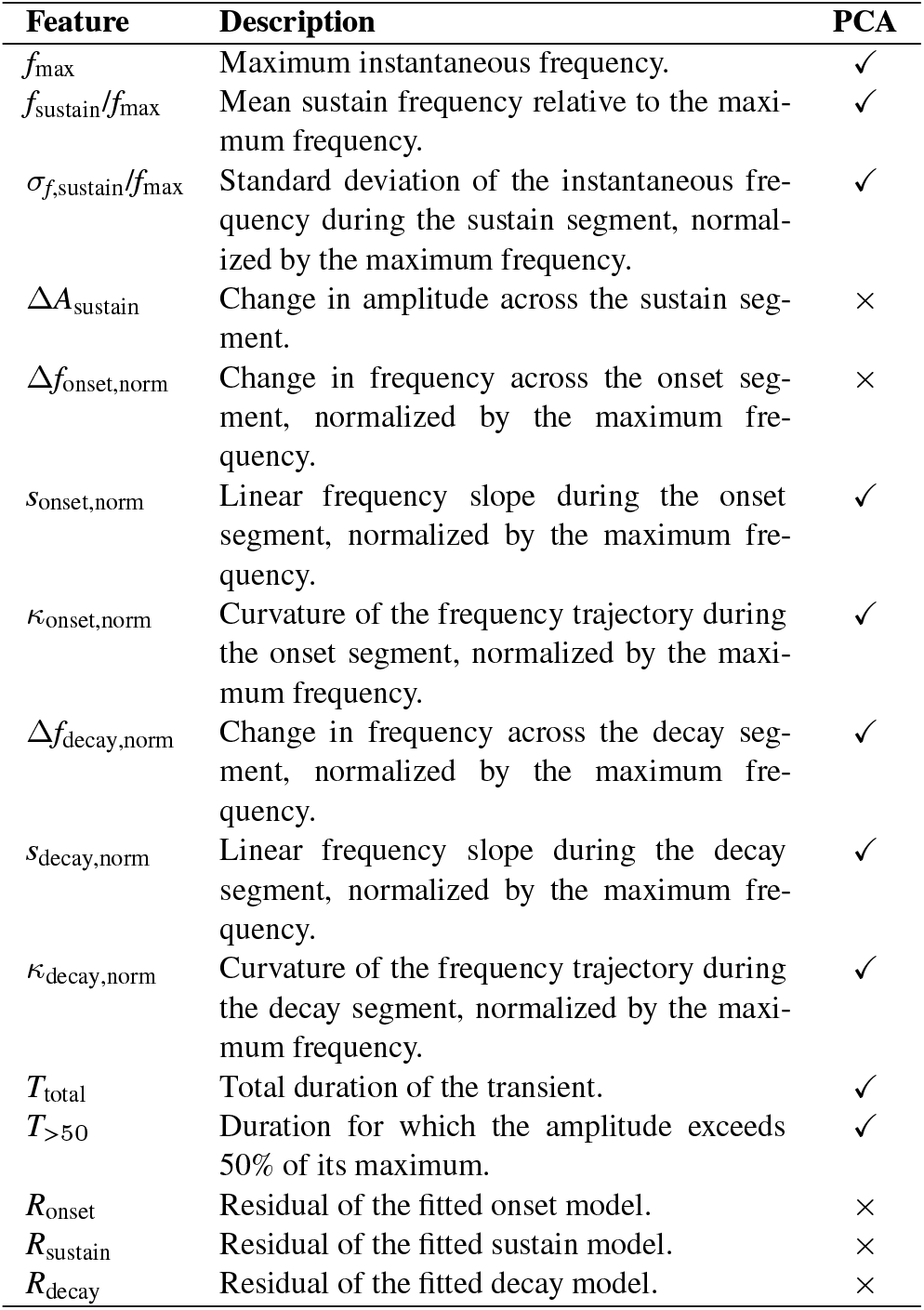
Features extracted from the instantaneous-frequency trajectory and their selection for the final PCA. Check marks indicate retained features and crosses indicate excluded features.

| Feature | Description | PCA |
| --- | --- | --- |
| $f_{\max}$ | Maximum instantaneous frequency. | ✓ |
| $f_{\text{sustain}}/f_{\max}$ | Mean sustain frequency relative to the maximum frequency. | ✓ |
| $\sigma_{f,\text{sustain}}/f_{\max}$ | Standard deviation of the instantaneous frequency during the sustain segment, normalized by the maximum frequency. | ✓ |
| $\Delta A_{\text{sustain}}$ | Change in amplitude across the sustain segment. | × |
| $\Delta f_{\text{onset},\text{norm}}$ | Change in frequency across the onset segment, normalized by the maximum frequency. | × |
| $s_{\text{onset},\text{norm}}$ | Linear frequency slope during the onset segment, normalized by the maximum frequency. | ✓ |
| $\kappa_{\text{onset},\text{norm}}$ | Curvature of the frequency trajectory during the onset segment, normalized by the maximum frequency. | ✓ |
| $\Delta f_{\text{decay},\text{norm}}$ | Change in frequency across the decay segment, normalized by the maximum frequency. | ✓ |
| $s_{\text{decay},\text{norm}}$ | Linear frequency slope during the decay segment, normalized by the maximum frequency. | ✓ |
| $\kappa_{\text{decay},\text{norm}}$ | Curvature of the frequency trajectory during the decay segment, normalized by the maximum frequency. | ✓ |
| $T_{\text{total}}$ | Total duration of the transient. | ✓ |
| $T_{>50}$ | Duration for which the amplitude exceeds 50% of its maximum. | ✓ |
| $R_{\text{onset}}$ | Residual of the fitted onset model. | × |
| $R_{\text{sustain}}$ | Residual of the fitted sustain model. | × |
| $R_{\text{decay}}$ | Residual of the fitted decay model. | × |

Figure 1 shows the amplitude–frequency trajectory of a European Stonechat (*Saxicola rubicola*) call. The inset displays the derived phase-based frequency trajectory overlaid on the spectrogram. Second-order polynomial regressions, displayed as blue and orange dashed lines, closely approximate the onset and decay segments, respectively.

**FIGURE 1.**
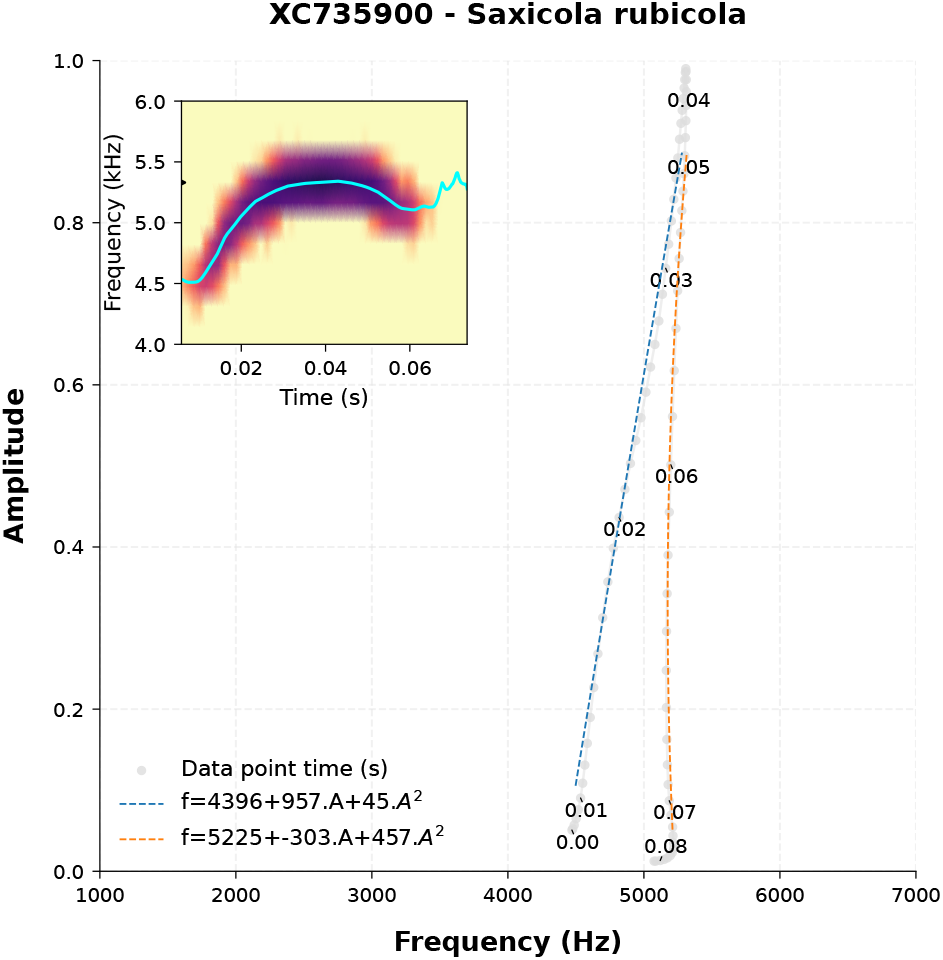
The amplitude-frequency trajectory of a European Stonechat call. Along the trajectory time is annotated in seconds. The inset shows the spectrogram overlaid by the derived time-frequency trajectory. The blue and orange dashed lines are the polynomial regressions to the onset and decay segments.

### 2.3 Dataset and call selection

We performed PCA on vocalisation data from species in the genus *Saxicola*, obtained from the Xeno-Canto website. These species produce two types of alarm calls: a broadband ‘chack’ and a narrowband, modulated ‘whit’. The analysis was restricted to the whit call type. Recordings with the highest or second-highest data quality ratings (‘a’ or ‘b’) were selected, provided they contained at least six calls. Metadata associated with each recording included location, recordist, date, and data quality. Table C.1 summarises the species and geographic origin of the recordings included in the analysis. The dataset had a geographic emphasis on Eurasian species, although several African species were also included.

Each call was extracted from the recording using a threshold set at 3% of the peak amplitude. In a handful of cases, low-frequency noise necessitated the application of a 3 kHz high-pass filter to ensure reliable onset detection.

When calls in a recording varied markedly in amplitude, implying possible different sources, those with peak waveform amplitude more than 6 dB below the maximum were marked by adding B to the recording name. The largest amplitude difference in the dataset occurred in recording XC594901, where the maximum spectrogram amplitudes of calls differed by 50 dB. This example illustrates the ability of the method to recover amplitude–frequency trajectories across a large range of signal levels.

Fewer than 10% of the records or calls within records were identified as clear outliers. Because outliers can substantially influence PCA, both conventional and robust PCA were performed.

### 2.4 Principal component analysis and robust PCA

Principal component analysis (PCA) was applied to the resulting feature vectors to identify the dominant dimensions of variation among bird vocalisations. Before analysis, the feature variables were standardised to zero mean and unit variance. PCA was performed on different combinations of taxonomic and geographic groups to investigate similarities and differences in their acoustic properties.

Both conventional PCA and robust PCA using Principal Component Pursuit (PCP) were performed. PCP decomposes the data matrix into low-rank and sparse components. The decomposition was computed using the Alternating Direction Method of Multipliers, following Candès et al. (2011).

### 2.5 PCA visualisation and individual separation

To visualize the distribution of calls among individuals, the calls were displayed in the space defined by three selected principal components. Calls belonging to the same individual were plotted as a set, with each individual represented by a centre of gravity (CoG), calculated as the mean position of its calls in the three-dimensional PCA space. The calls were connected to their respective CoG by faint lines to show both the location and dispersion of each call set. CoGs were annotated with a unique numerical identifier corresponding to the individual, while colour was used to distinguish species. This representation therefore allows both within-individual variation and between-individual separation to be assessed simultaneously.

For individual *i*, with *n*_*i*_ calls represented by the three selected principal components, the CoG was calculated as

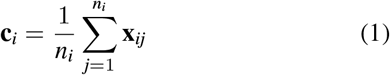

where (**x**_*ij*_) denotes the three-dimensional PCA coordinate of call *j* from individual *i*. The lines connecting individual calls to their CoG provide a visual representation of within-individual dispersion. This dispersion was quantified as the mean Euclidean distance of the calls from their CoG,

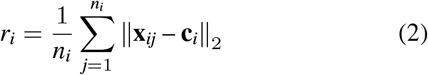

To quantify the separation of each individual from the other individuals, the distance 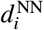 neighbour, was calculated as from its CoG to the nearest neighbour, was calculated as

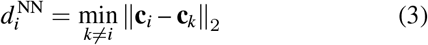

The relative separation of an individual was then expressed as the ratio

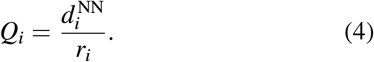

A large *Q*_*i*_ indicates that an individual’s CoG is relatively distant from its nearest neighbouring individual compared with the spread of its own calls, whereas a small *Q*_*i*_ indicates that the individual’s call set is relatively close to another individual compared with its within-individual dispersion.

The CoG marker sizes in the plots were scaled according to *Q*_*i*_. To reduce the influence of extreme values, the 10th and 90th percentiles of the *Q*_*i*_ distribution, denoted by *Q*_10_ and *Q*_90_, respectively, were used as the lower and upper limits of the scaling range. The marker size was calculated as

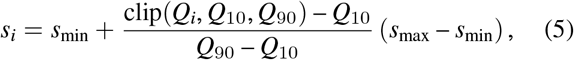

where *s*_min_ and *s*_max_ are the minimum and maximum marker sizes. Values of *Q*_*i*_ below *Q*_10_ were therefore assigned the minimum marker size, and values above *Q*_90_ the maximum marker size. Consequently, larger CoG markers identify individuals whose call sets are relatively well separated from their nearest neighbours, taking within-individual variation into account.

## 3 PCA STRUCTURE AND STATISTICAL INTERPRETATION

### 3.1 Conventional and robust PCA

Conventional principal component analysis (PCA) is sensitive to outliers because it is based on squared deviations from the mean, allowing extreme observations to exert a disproportionate influence on the resulting principal components. Robust PCA using Principal Component Pursuit (PCP) addresses this limitation by decomposing the feature matrix into a low-rank component, representing the underlying structure of the data, and a sparse component, capturing unusual observations or feature values. The decomposition uses the *L*_1_ norm to penalise the sparse component. Unlike a squared penalty, this penalty increases linearly with the magnitude of a deviation, reducing the disproportionate influence that large deviations can have on the decomposition. When a dataset contains few influential outliers, conventional and robust PCA may yield similar results, although their agreement depends on the structure of the data.

### 3.2 Explained variance and feature loadings

The effectiveness of robust PCA for the *Saxicola* dataset can be assessed by comparing its loading matrix with that obtained using conventional PCA. The analysis used the complete dataset, comprising 801 calls from 49 recordings representing 15 species. Figure 2 shows the conventional PCA loadings, while Figure 3 shows the robust PCA loadings. Each cell represents the loading of a feature on a principal component, indicating the direction and strength of their association. The proportion of variance explained by each component is given below the corresponding column.

**FIGURE 2.**
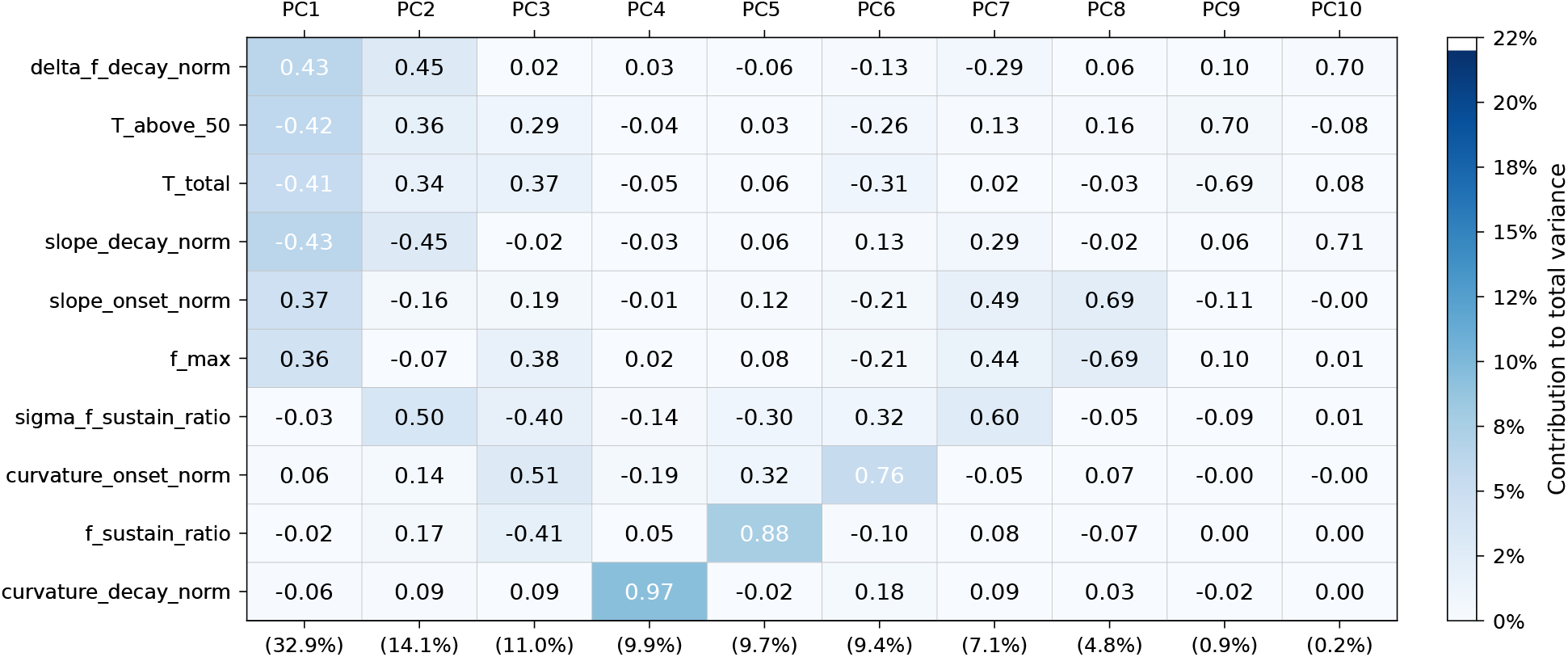
The loading matrix for the PCA. The number in each cell gives the contribution, or loading, of the features to the principal components. The colour of each cell represents the contribution of the total variance of each feature, weighted by the overall contribution of that principal component.

**FIGURE 3.**
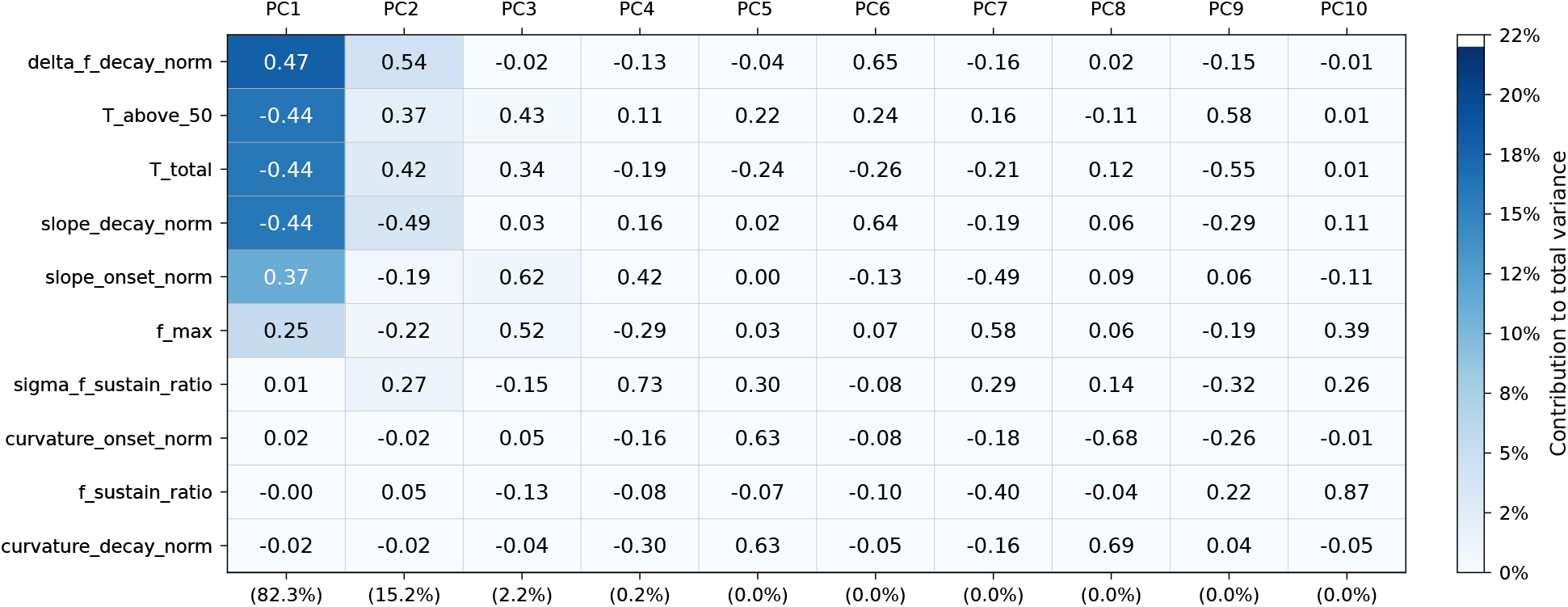
The loading matrix for the Robust PCA

Conventional PCA exhibits a relatively flat explained-variance curve, requiring nine of the ten components to account for more than 99% of the variance in the original feature matrix. In contrast, applying PCA to the low-rank component obtained through robust PCA achieves the same level of explained variance with just three components. This indicates that, for this dataset, the robust low-rank representation concentrates the structured variation into fewer dimensions, providing a more compact representation for subsequent analysis. This is an empirical result and does not imply that robust PCA will always yield a more compact representation.

The cell colours represent each feature’s contribution to the total dataset variance, calculated by multiplying its proportionate contribution to a principal component, based on its squared loading, by the proportion of variance explained by that component. The relatively light colours associated with PC3 indicate that it accounts for only a small proportion of the total dataset variance. However, some features have substantial loadings on PC3, indicating an association with variation along this axis despite its relatively small contribution to overall variance. The numerical value in each cell gives the signed loading, indicating the direction and strength of the feature’s association with the component.

Based on the feature loadings, the three components can be interpreted approximately as follows:

- **PC1:** dominant frequency and frequency drift.
- **PC2:** temporal and decay characteristics.
- **PC3:** curvature and sustain structure.

The dominant dimension identified by robust PCA was strongly correlated with the corresponding dimension from conventional PCA (*r* = 0.95). However, PC1 accounted for 82.3% of the variance in the robust low-rank representation, compared with 32.9% of the variance in the original feature matrix. These percentages refer to different matrices and therefore should not be interpreted as a direct comparison of the proportion of original dataset variance explained by the two methods. The dominant dimension is primarily associated with features describing the frequency trajectory of the transient, particularly the maximum frequency, the normalised decay-frequency change, and the onset and decay slopes and curvatures.

### 3.3 Interpretation of PCA structure and individual separation

The difference between the multi-species and single-species PCA representations reflects how PCA allocates its principal axes to the dominant sources of variation. When multiple species are included, between-species differences contribute substantially to the total covariance and therefore occupy prominent directions in PCA space. The leading principal components consequently capture a combination of species-level and within-species variation.

When the analysis is restricted to a single species, this major source of between-species variation is removed. The PCA axes are consequently reoriented to represent the variation that remains, which can include differences among individuals and calls within the species. The apparent increase in separation between individual call sets in the single-species representation therefore does not necessarily indicate an increase in the underlying acoustic differences between individuals. Rather, the coordinate system has changed so that within-species variation becomes more prominent in the resulting representation.

This distinction is important because the first three principal components account for almost all of the variance in both analyses, even though they represent different sources of variation. The increased separation of individuals in the single-species representation is therefore not simply a consequence of retaining less information in the three-dimensional display. Instead, the PCA axes capture different aspects of the acoustic variation, with the multi-species analysis reflecting substantial species-level structure and the single-species analysis potentially emphasising within-species and individual-level differences.

The distribution of individual call sets and their centres of gravity (CoGs) provides a further means of examining this effect. The separation score (*Q*_*i*_), defined as the distance from an individual’s CoG to its nearest neighbouring CoG relative to the mean dispersion of its own calls, quantifies the separation of individual call sets in relation to their within-individual variation. The single-species PCA does not create additional acoustic information; rather, it changes the relative prominence of the variation represented by the selected axes.

Some structure was also visible when the calls were projected onto higher principal components (PC4–PC6), although these components accounted for very little of the total variance. In the present analysis, PC4 explained only 0.73% of the variance and PC5 explained 0.12%, while the remaining components contributed negligible amounts. Individual separation was substantially lower in the PC4–PC6 space than in the PC1–PC3 space, with median *Q*_*i*_ values of 0.21 and 1.01, respectively. Thus, the structure visible in the higher principal components did not correspond to greater separation of individual call sets relative to their within-individual dispersion. Although higher components can reveal additional patterns of variation, these results indicate that the structure observed in PC4–PC6 provides a weaker representation of individual separation in this dataset.

The stability of the PCA representation when new calls are introduced was assessed by incrementally adding the 15 calls from recording XC1105030 to a reference dataset of 789 calls representing 15 species. The identification of XC1105030 as *S. torquatus* is flagged as “under discussion” on Xeno-Canto. After all 15 calls had been added, the angles between the original and updated principal-component axes were 0.49°, 0.67°, and 1.67° for PC1, PC2, and PC3, respectively. These small angular changes indicate that the principal-component directions remained largely stable under this particular addition, especially for the first two components. Figure 4 shows the cumulative angular changes as successive calls were added. Thus, the inclusion of the disputed recording had little effect on the dominant acoustic structure captured by the robust PCA representation.

**FIGURE 4.**
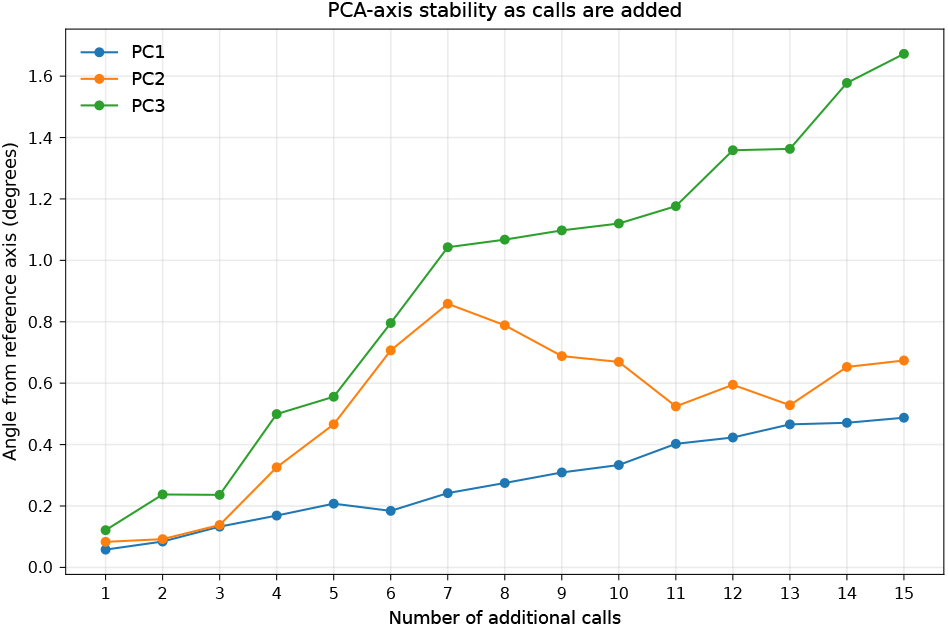
Stability of the PCA orientation as successive calls are added.

To examine the acoustic relationship between the disputed recording and recordings with existing species assignments, we examined its position in the robust PCA representation fitted to the dataset including these calls. Figure 5 shows that the calls from XC1105030 (recording 49, red box), although assigned to *S. torquatus*, fell within the spread of calls assigned to *S. rubicola*. In contrast, the other recordings assigned to *S. torquatus* (recordings 21, 45, and 46, blue box) were well separated from the *S. rubicola* calls. Thus, the disputed recording exhibits an acoustic pattern that differs from those of the other *S. torquatus* recordings in the dataset and overlaps with the observed variation in *S. rubicola*. This result raises the possibility that its acoustic characteristics are more consistent with those of *S. rubicola*, but does not establish its taxonomic identity. The example illustrates how robust PCA can help identify recordings whose acoustic characteristics warrant further investigation.

**FIGURE 5.**
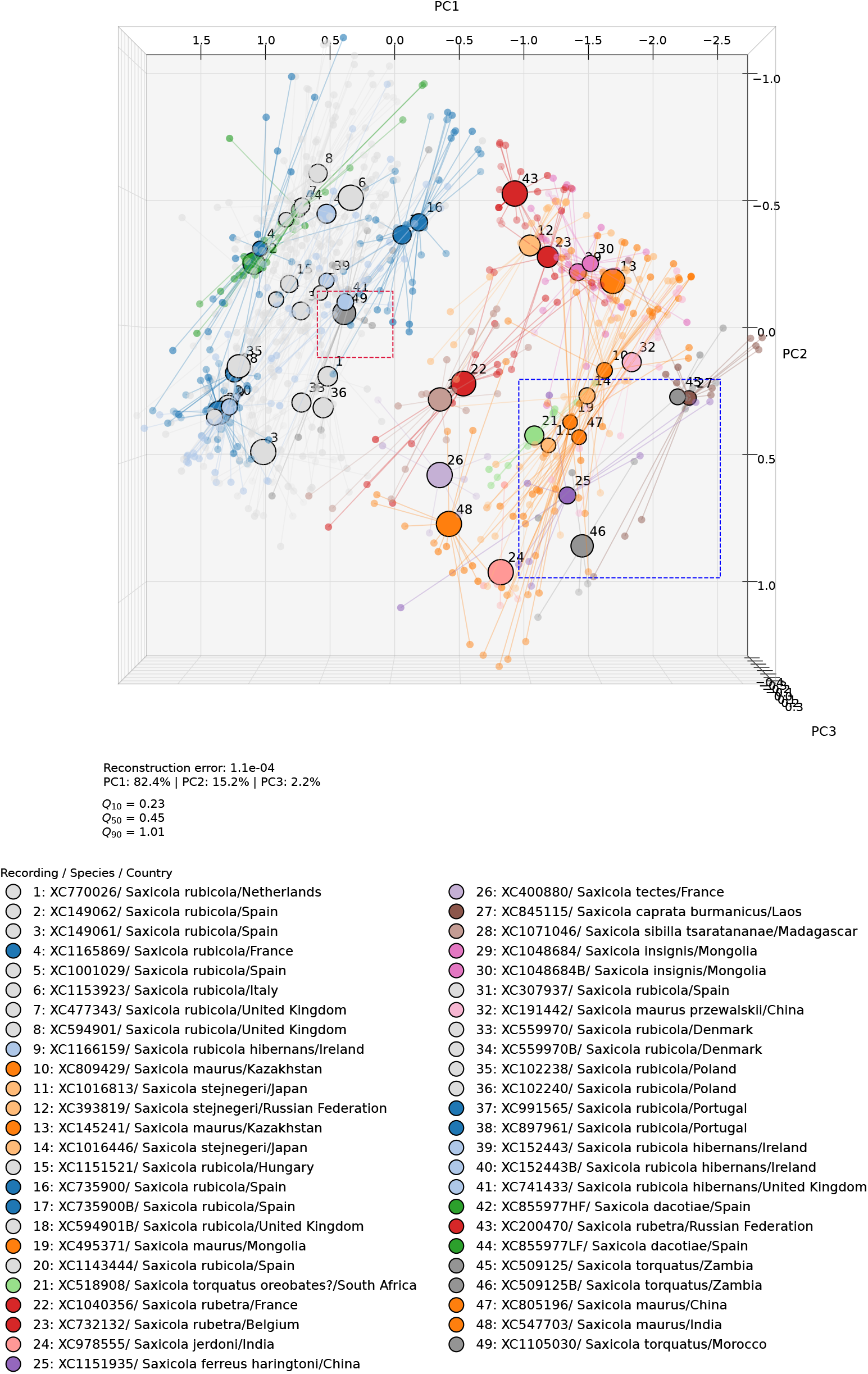
Robust PCA representation of the *Saxicola* dataset, showing the calls from a recording identified as *S. torquatus* whose identification is under discussion on Xeno-Canto. The disputed calls (no. 49) are highlighted with the red box. The blue box indicates the spread of the *S. torquatus* calls 21, 45 and 46.

Taken together, these analyses indicate that robust PCA provides a compact and relatively stable representation of the dominant acoustic variation in the *Saxicola* dataset. Its application to the disputed recording illustrates how this representation can reveal acoustic similarities and differences between recordings with different species assignments. Although PCA alone cannot establish taxonomic identity, it provides a useful framework for investigating uncertain identifications in the context of existing acoustic variation.

## 4 BIOLOGICAL INTERPRETATION OF SAXICOLA VOCALISATIONS

### 4.1 Interspecific variation

The taxonomy of the *Saxicola torquatus* complex illustrates the challenges of delimiting species and reconstructing their evolutionary relationships. Mitochondrial markers have provided evidence for distinct evolutionary lineages, but their taxonomic interpretation has remained debated. Opaev et al. (2018), for example, combined genetic evidence with morphology, plumage, song, and alarm-call characteristics to support the recognition of three northern Eurasian species: *S. rubicola, S. maurus* and *S. stejnegeri*. Their findings illustrate the potential value of vocal characteristics as complementary evidence in taxonomic assessment, particularly where other diagnostic features are limited.

Phylogenetic and biogeographic studies have also used *Saxicola* to investigate geographic diversification and speciation (Illera et al., 2008). Recent phylogenomic work further highlights the complexity of evolutionary relationships within the *S. torquatus* complex, reporting patterns consistent with introgression and incomplete lineage sorting (Jiang et al., 2025, 2026). Genetic studies differ in their sampling and sequencing strategies. For example, Jiang et al. (2026) report genomic data from a single S. stejnegeri individual, whereas their analyses also incorporate pooled sequencing data from 51 S. torquatus and 56 S. dacotiae individuals. Pooled sequencing can capture genetic variation across multiple individuals, but does not retain the same individual-level information as separately sequenced samples. Quantitative acoustic analysis offers a complementary approach for characterising vocal variation across calls, individuals and geographic locations, provided that individual identity and sampling structure are taken into account.

Quantitative acoustic characterisation could contribute to investigations of species boundaries and geographic differentiation by enabling systematic comparisons of vocal variation across species, populations and regions. When individual identity and sampling structure are taken into account, such comparisons can help distinguish variation within individuals and populations from differences among them. Combined with genetic, morphological and distributional evidence, acoustic analyses could therefore help evaluate competing taxonomic and biogeographic hypotheses. The present study does not address these evolutionary questions directly, but demonstrates an approach that could support such investigations.

The patterns in Figure 6 illustrate the potential of this approach for examining interspecific acoustic variation. In the selected PCA representation, calls assigned to *S. rubicola*, including the subspecies *S. r. rubicola* and *S. r. hibernans*, are separated from those assigned to *S. rubetra, S. insignis* and *S. maurus*. The centres of gravity for *S. dacotiae* occupy positions that can be compared with those of the other taxa in this acoustic feature space. Such patterns identify similarities and differences in the measured vocal characteristics, but should not in themselves be interpreted as evidence of phylogenetic relationships.

**FIGURE 6.**
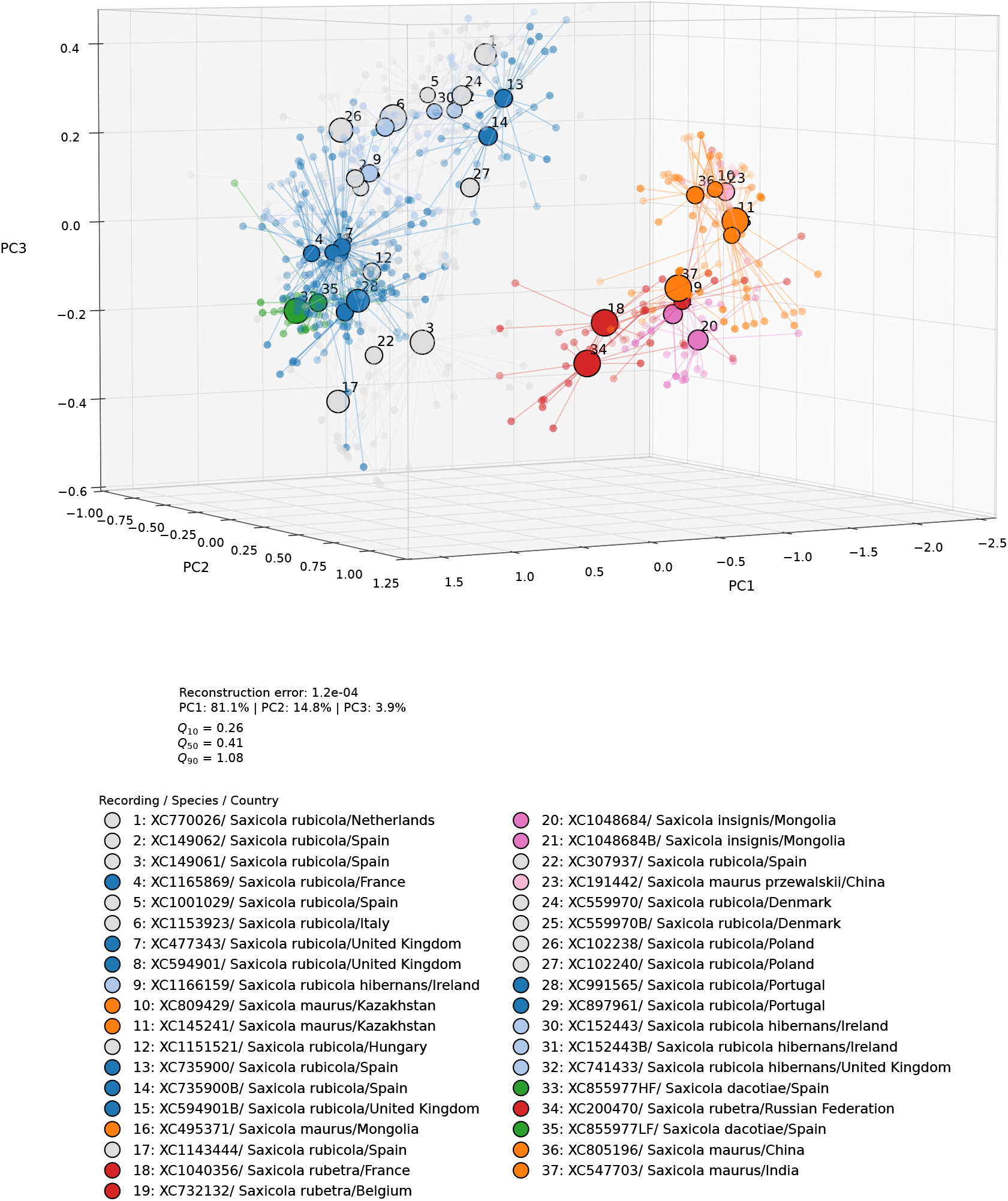
Three-dimensional PCA representation of interspecific variation in vocalisations of European and Asian species from the *Saxicola* genus.

### 4.2 Intraspecific and individual variation

Variation within *Saxicola rubicola* provides an opportunity to investigate whether quantitative acoustic characteristics can help distinguish differences between subspecies from variation among individuals. However, such comparisons are complicated by uncertainty in subspecific identification. Urquhart and Bowley (2002) noted that although *S. r. hibernans* is marginally paler than *S. r. rubicola*, distinguishing the two sub-species where they meet is virtually impossible in the field. The reliability of labels assigned to recordings is therefore an important consideration when interpreting apparent acoustic differences between subspecies.

Figure 7 distinguishes *S. rubicola* calls recorded at coastal locations (dark blue) from those recorded inland (dark grey). Calls labelled as *S. r. hibernans* in Xeno-Canto are shown in light blue. The PCA representation shows no clear separation between these groups based on the acoustic features analysed. This overlap may reflect genuine similarity in vocal characteristics, individual variation, geographic variation, or uncertainty in the taxonomic labels; the present data do not allow these possibilities to be distinguished.

**FIGURE 7.**
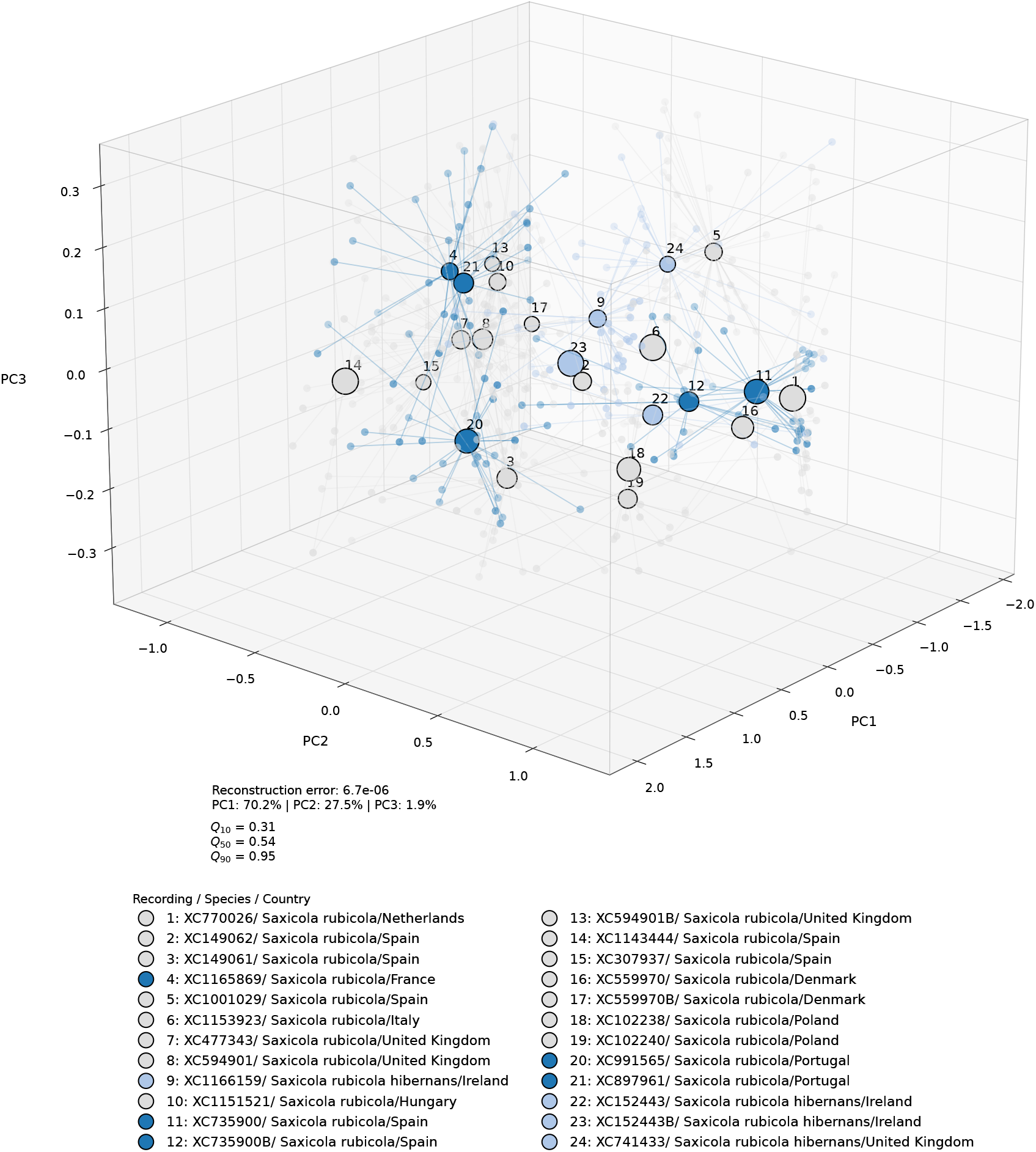
PCA representation of calls assigned to *S. r. rubicola* from two recordings made at different locations along the Portuguese coast. Calls from a second individual recorded simultaneously are identified by the suffix ‘B’ appended to the recording identifier.

The identity of stonechats breeding in Portugal provides a relevant example. Siddle (2006), in discussing which sub-species of Common Stonechat breeds in Portugal, concluded that the birds were more similar to *S. r. rubicola* than to a clinal extreme of *S. r. hibernans*, based on his observations and published descriptions. To explore this question acoustically, we compared calls from two recordings made at different locations along the Portuguese coast during the breeding season with calls assigned to *S. r. rubicola* in the PCA representation (Figure 7, recordings 20 and 21). Their positions relative to the reference calls provide a preliminary indication of acoustic similarity to *S. r. rubicola*. However, the small number of recordings precludes firm conclusions about the subspecific identity of stonechats in Portugal.

The same PCA representation includes calls from recordings made in Ireland (recordings 22 and 23, involving a male and female, with juveniles probably nearby), Denmark (recording 16, a female, and recording 17, a male), the United Kingdom (a nearby male in recording 8 and a more distant female in recording 13), and Spain (recordings 11 and 12, involving a male and a juvenile female approximately 3 m apart). Field observations indicate that more than one individual was present in each case. All calls were normalised to the same amplitude before feature extraction, allowing their positions in PCA space to be compared independently of differences in overall recorded amplitude.

Except for the Danish pair, the paired calls remain close to one another in PCA space. The greater separation of the Danish pair indicates a larger difference in the selected acoustic features. Given the field observations that a female and a male were present, one tentative possibility is that the calls represent individuals from different social groups, or that one bird was outside the family group represented by the other. However, the available evidence does not establish the identity of the call producers or their social relationships. These examples illustrate how quantitative acoustic analysis may reveal differences between calls that warrant further investigation at the individual and social-group levels.

### 4.3 Geographic variation

The PCA representation reveals three broad geographic groupings of calls (Figure 8). Group A comprises recordings from Europe. Group B comprises recordings from the non-European Palearctic, although it also includes European recordings of the Whinchat (*Saxicola rubetra*). Group C comprises recordings from the Afrotropical and Indomalayan realms, including Africa, Madagascar, Réunion, Laos and southwestern China. These groupings describe geographic patterns in acoustic feature space rather than formally defined biogeographic regions, and their interpretation must account for overlap between groups and the geographic distribution of the sampled species. The three-dimensional representation in Figure 6 shows a substantial separation between groups A and B. In contrast, Figure 9 shows a more gradual transition from species in group B towards the African species in group C, with South Asian species occupying intermediate positions in acoustic feature space. This pattern suggests that the acoustic differences between these groups are not uniformly discrete. However, the extent to which the apparent continuum reflects geographic variation, differences between species, or both requires further investigation.

**FIGURE 8.**
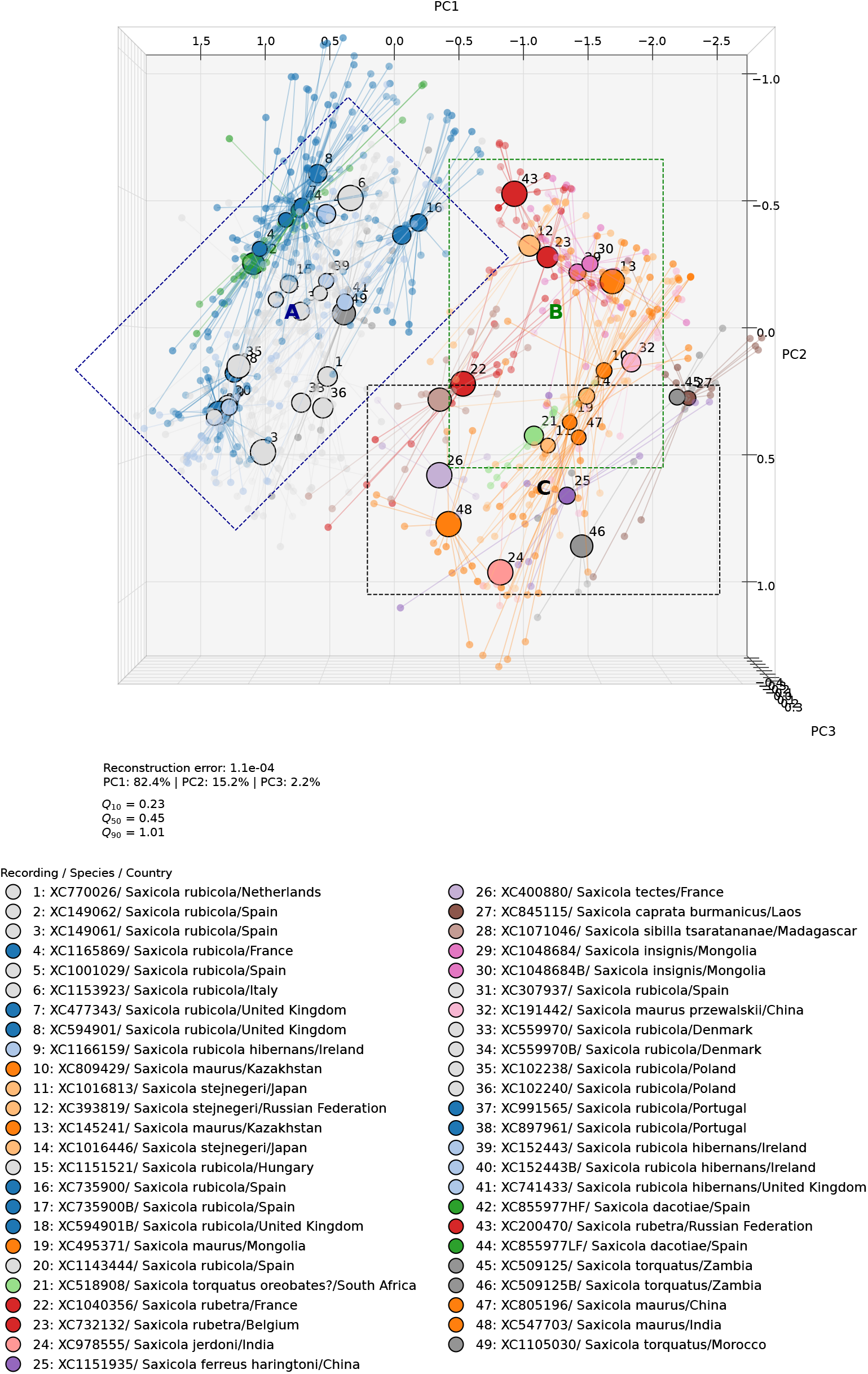
Robust PCA representation of the *Saxicola* dataset. Group A comprises recordings from Europe, Group B comprises recordings from the non-European Palearctic, and Group C comprises recordings from the Afrotropical and Indomalayan realms. Recordings of the Whinchat (*S. rubetra*) made in Europe are assigned to Group B.

**FIGURE 9.**
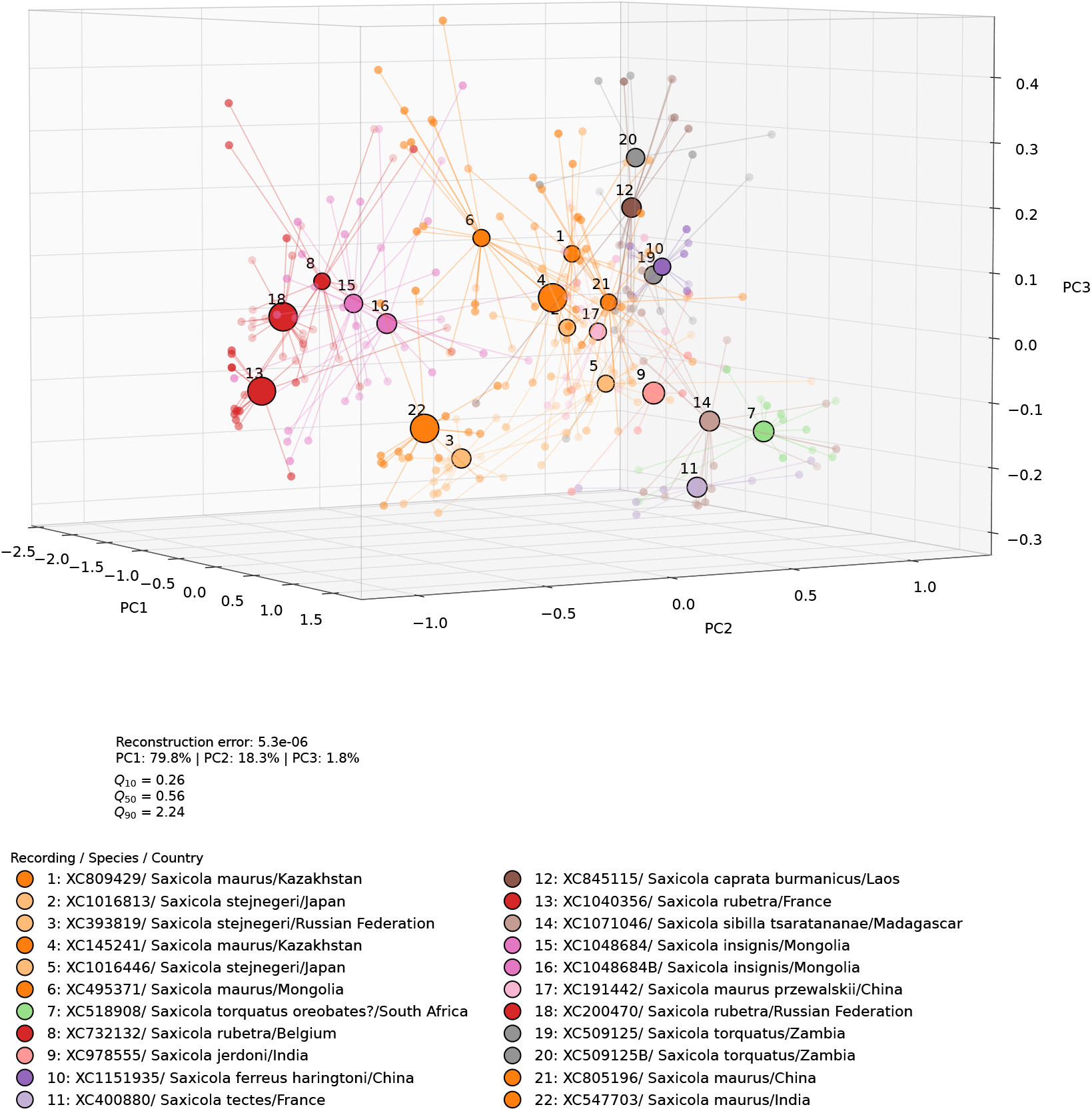
Three-dimensional PCA representation illustrating the gradual transition in acoustic space from Group B towards Group C.

The ordering of species in acoustic feature space may also help generate hypotheses about evolutionary relationships and potential gene flow. For example, the proximity of *S. maurus* and *S. stejnegeri* is consistent with the hypothesis of gene flow between these taxa, discussed by Jiang et al. (2026), and motivates further investigation of the relationship between acoustic similarity and introgression. More broadly, the positions of species in acoustic space provide an independent perspective that can be compared with proposed phylogenetic and biogeographic relationships. However, acoustic similarity alone cannot establish evolutionary relationships, hybridisation or gene flow, nor does acoustic dissimilarity necessarily contradict a shared ancestry. Combining quantitative acoustic comparisons with genomic data and geographically explicit sampling could help investigate how vocal variation relates to genetic differentiation and evolutionary history.

The vocalisations of *Saxicola dacotiae*, which is generally considered genetically closely related to *S. rubicola*, exhibit a distinctive acoustic structure. All *S. dacotiae* calls examined in the Xeno-Canto recordings show two simultaneous, closely spaced spectral components at approximately 6–7 kHz. The frequency separation observed for the XC855977 recording corresponds to an amplitude-modulation rate of approximately 776 Hz, producing a characteristic beat pattern visible in Figure 10. To examine the contributions of the two components separately, we applied low-pass and high-pass filters to obtain the lower- and higher-frequency components, respectively. Figure 11 shows the high-pass-filtered component. In the PCA plots, these components are identified as XC855977LP and XC855977HP, respectively.

**FIGURE 10.**
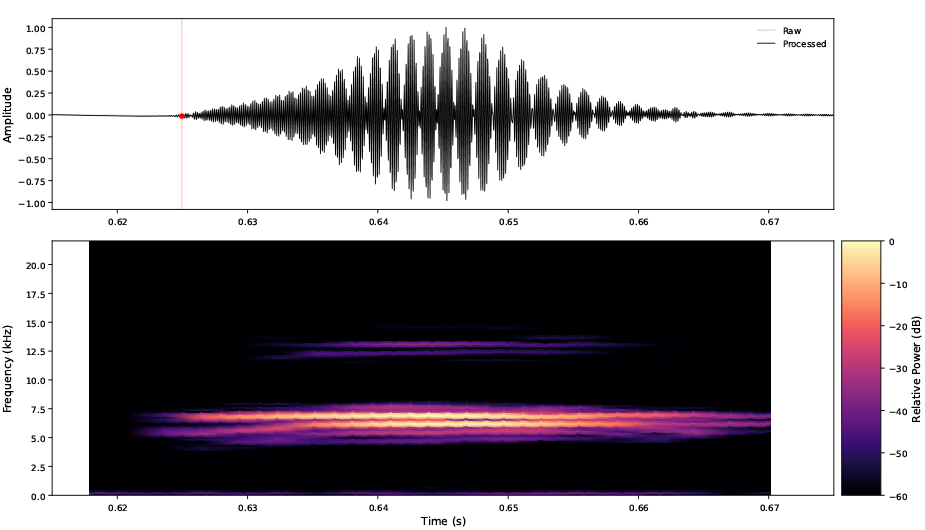
The vocalisations of *Saxicola dacotiae* exhibit a distinctive acoustic structure. The calls show two simultaneous, closely spaced spectral components at approximately 6–7 kHz, resulting in a beat pattern for the calls of recording 5977 of 776 Hz.

**FIGURE 11.**
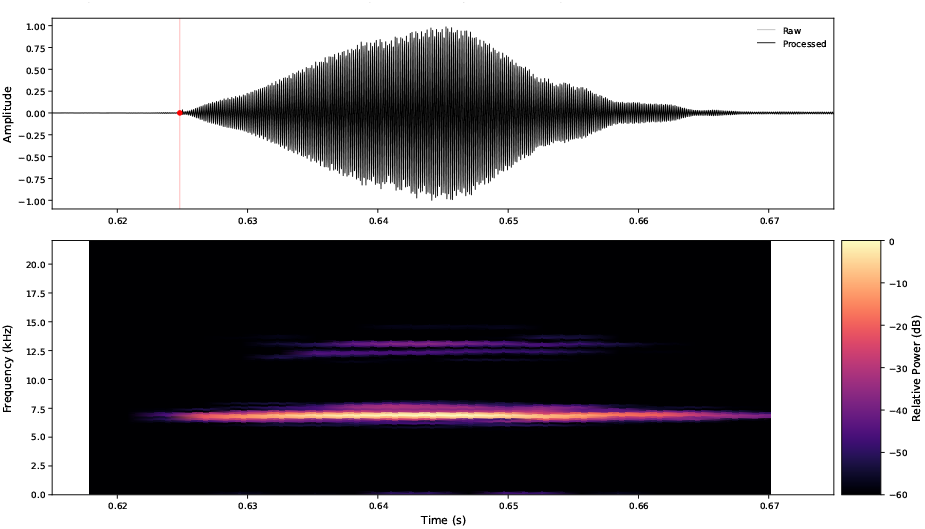
A FIR filter isolates the high-pass component of the *Saxicola dacotiae* calls from recording XC855977 (Figure 10). This reduces the beating pattern, allowing the call to be segmented into onset, sustain, and decay phases.

**FIGURE 12.**
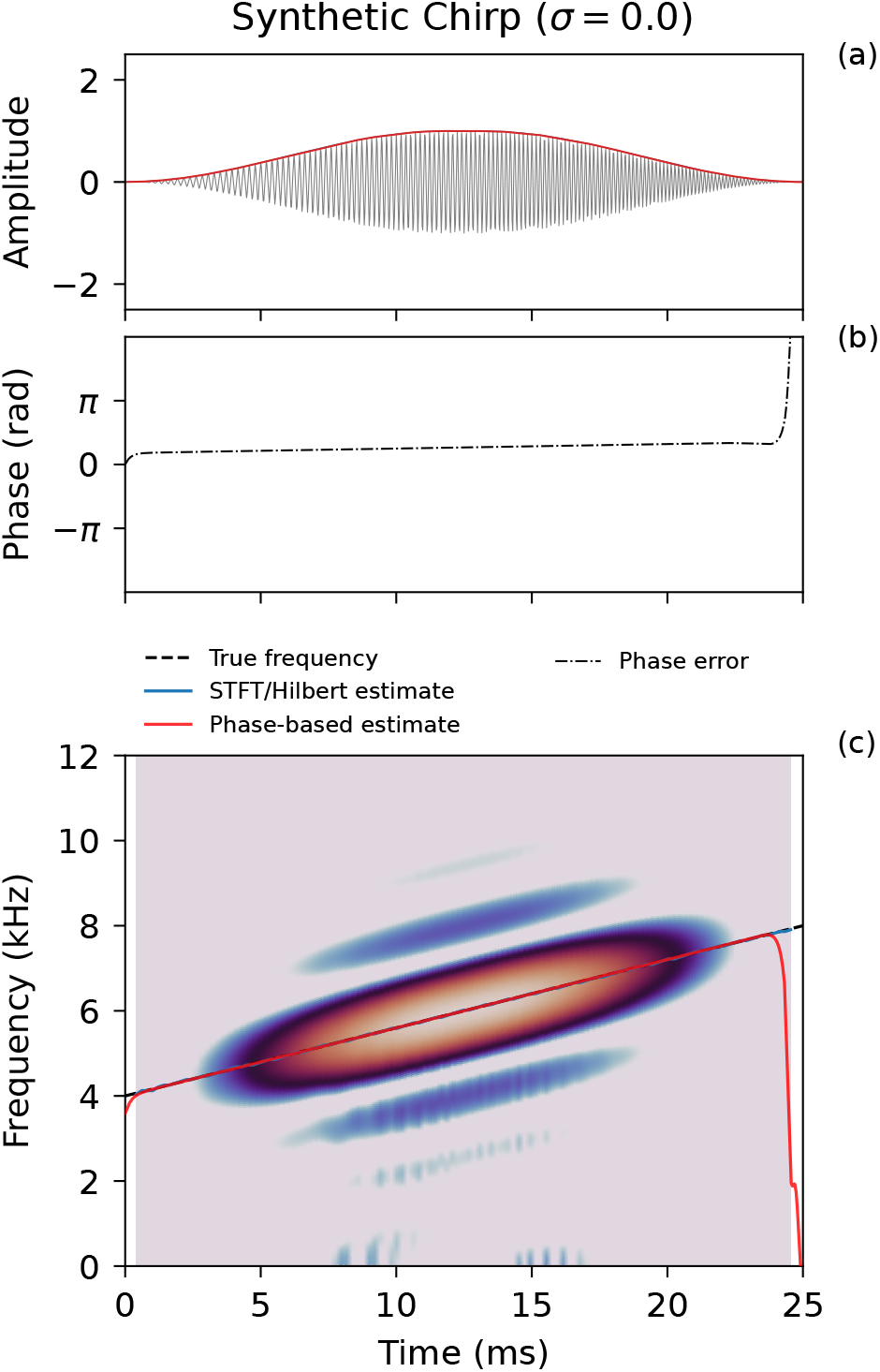
Tapered synthetic chirp: (a) Time. (b) Difference between theoretical phase and estimated phase. (c) Spectrogram showing the theoretical frequency (black dashed line), the STFT estimate (blue line), and the phase-based estimate (red line).

**FIGURE 13.**
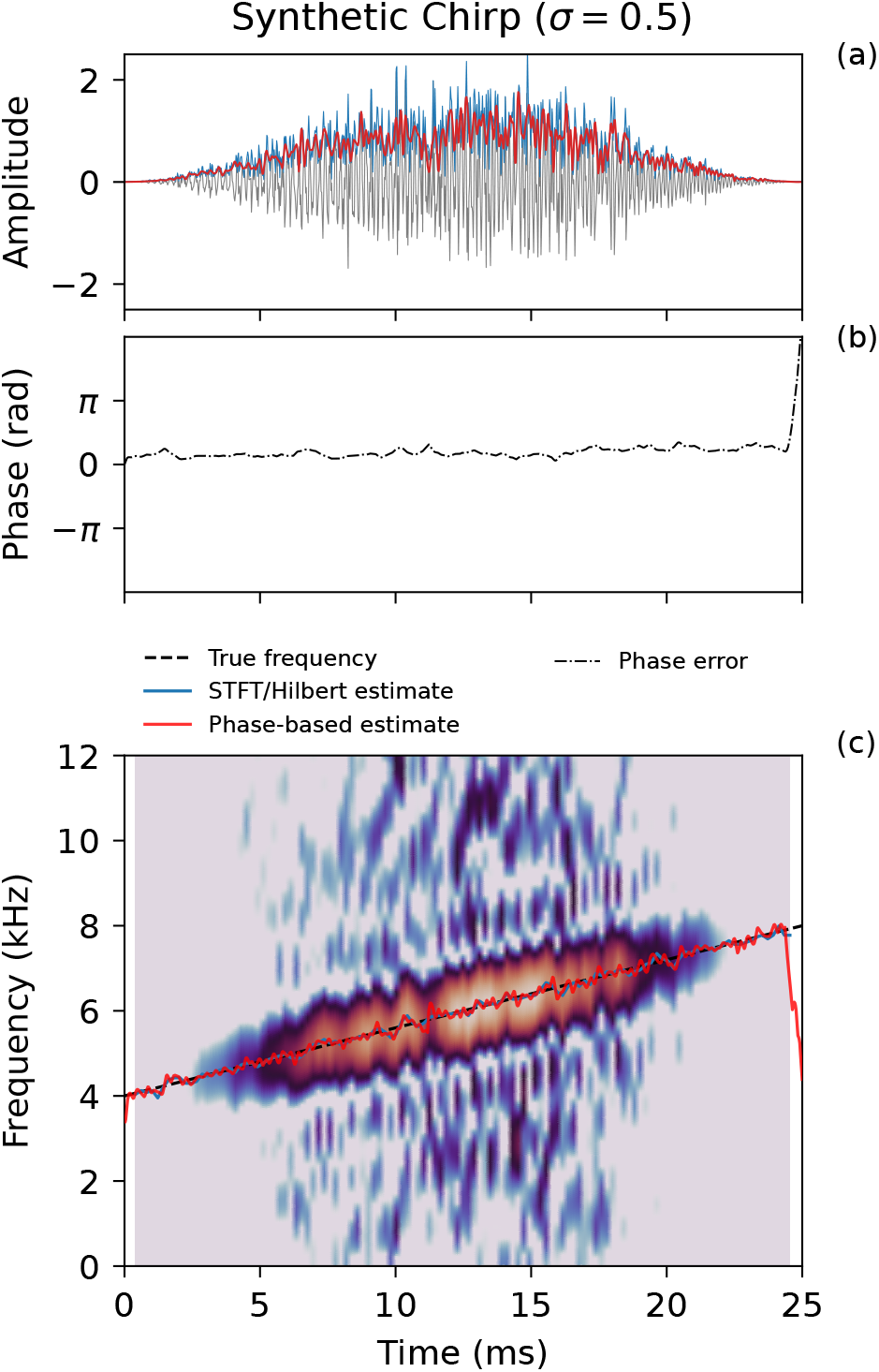
As in Figure 12, but with added 0.5 standard deviation noise.

**FIGURE 14.**
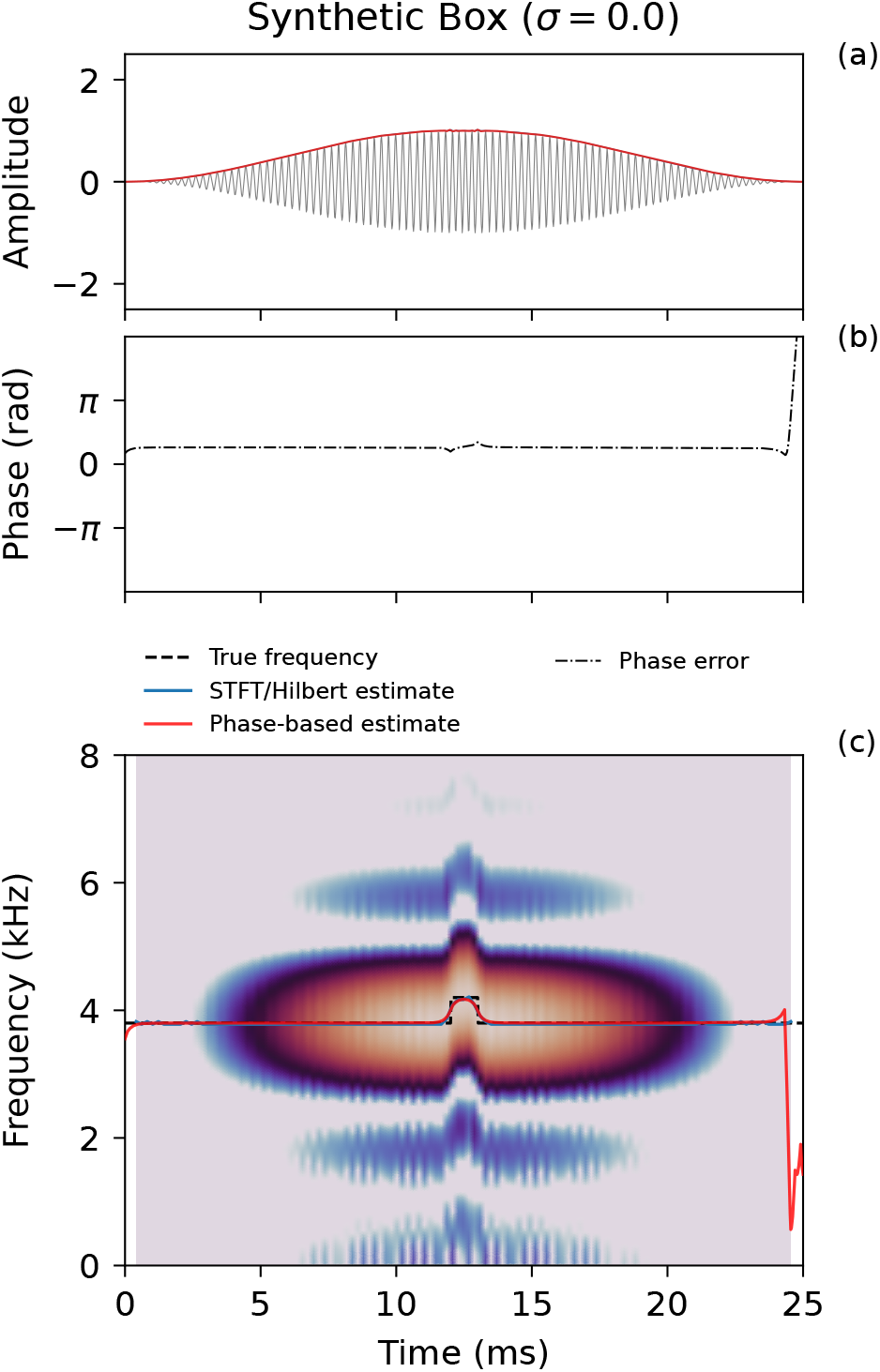
Tapered constant-frequency synthetic with a box-shaped 400 Hz frequency jump of 1-millisecond duration. (a) Time. (b) Difference between theoretical phase and estimated phase. (c) Spectrogram showing the theoretical frequency (black dashed line), the STFT estimate (blue line), and the phase-based estimate (red line).

**FIGURE 15.**
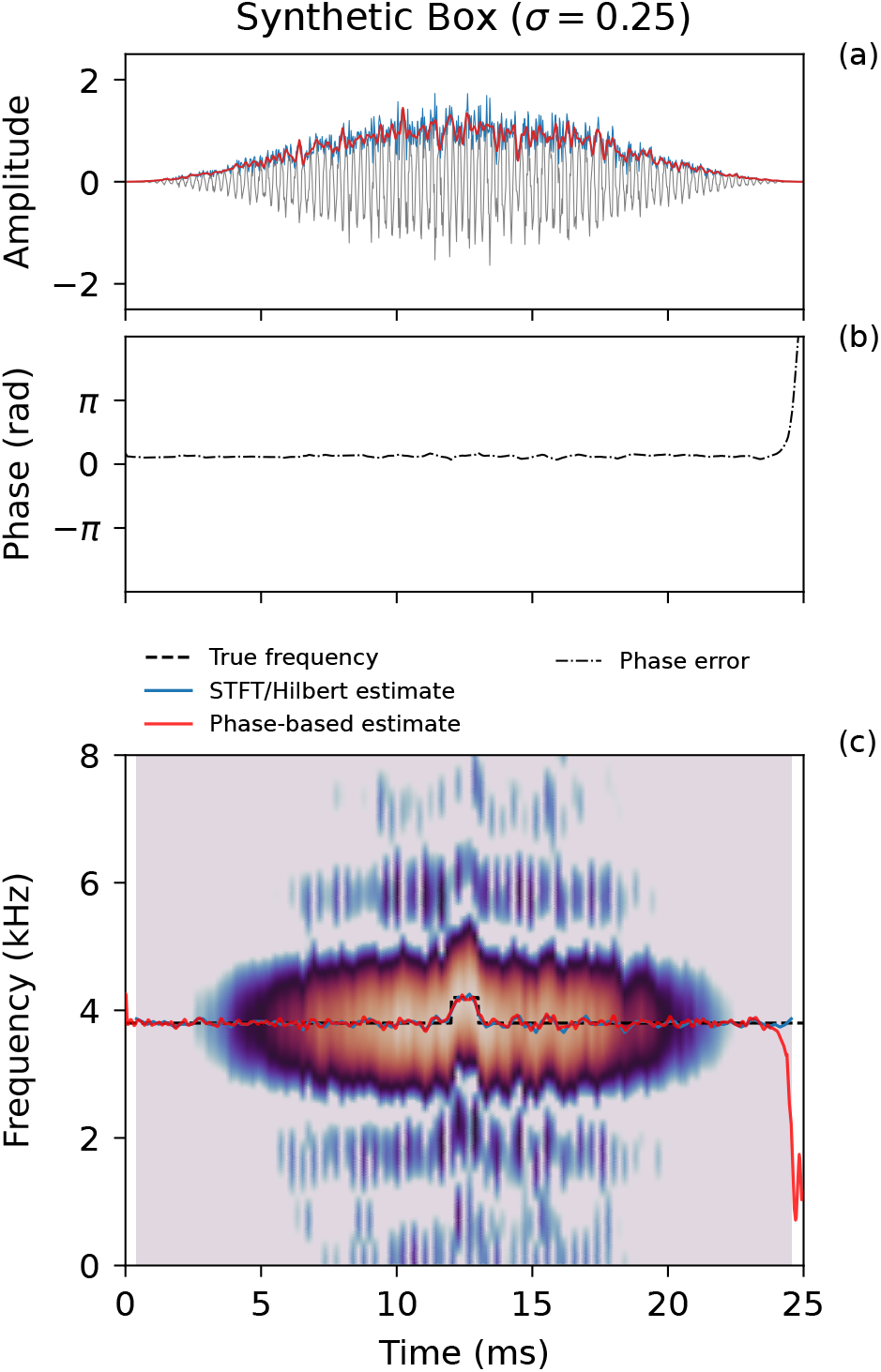
As in Figure 14, but with added 0.25 standard deviation noise.

## 5 DISCUSSION

This study demonstrates the feasibility of phase-based frequency tracking for the quantitative characterisation of alarm calls in the genus *Saxicola*. By representing amplitude as a function of instantaneous frequency, the method provides a basis for extracting acoustic features that describe the frequency and temporal structure of individual calls. Combined with principal component analysis (PCA), these features allow calls to be compared within a common acoustic representation, revealing patterns of similarity and variation across the sampled taxa. The results establish the potential of this approach for comparative acoustic analysis, while also defining the scope within which the present implementation has been evaluated.

The segmentation of calls into onset, sustain and decay phases exploits the relatively structured frequency trajectories and predominantly monochromatic character of the *Saxicola* calls analysed. This makes it possible to define and extract features associated with distinct parts of a call. The segmentation procedure and resulting feature set are therefore tailored to the vocal characteristics examined here and should not be assumed to transfer directly to other bird taxa. Applying the method to vocalisations with more complex frequency trajectories or multiple simultaneous spectral components would require appropriate adaptations to the segmentation and feature extraction procedures. This taxon-specific implementation does not, however, preclude the broader application of phase-based frequency tracking to other types of vocalisation.

The PCA results show that the extracted features capture substantial acoustic variation among the calls analysed. However, positions and separations in PCA space describe acoustic relationships within the selected feature representation; they do not directly establish taxonomic or evolutionary relationships. The observed patterns may reflect differences among species, geographic variation, individual variation, or a combination of these factors. Similarly, acoustic overlap does not necessarily imply taxonomic equivalence, and acoustic separation does not in itself establish distinct species status. The analyses are therefore best regarded as a means of characterising acoustic variation and identifying patterns that can be investigated further, rather than as a standalone basis for taxonomic classification.

The taxonomic and evolutionary questions surrounding several *Saxicola* taxa provide a relevant context for further development of this approach. Quantitative acoustic comparisons could complement genetic, morphological and distributional evidence by identifying similarities and differences that merit closer investigation. For example, acoustic relationships among taxa could be compared with proposed phylogenetic relationships or hypotheses of geographic differentiation and gene flow. Such comparisons would not establish hybridisation, introgression or shared evolutionary history from acoustic evidence alone, but could help formulate testable hypotheses for investigation using independent genetic data.

The dataset provides a substantial collection of calls for demonstrating the method, but its taxonomic and geographic coverage is uneven. Some species are represented by numerous calls, whereas others are represented by only a few recordings or lack sufficient high-quality alarm-call material for inclusion. Moreover, the number of calls analysed does not necessarily reflect the number of independent individuals sampled. Multiple calls from one individual can characterise within-individual vocal variation, but cannot substitute for sampling multiple individuals when assessing population-level or species-level differences. The interpretation of acoustic patterns must therefore take account of the available sample sizes, the distribution of recordings among taxa and locations, and uncertainty in individual identity and taxonomic labels.

The primary aim of this study is to demonstrate the feasibility of the method and its potential for comparative acoustic analysis, rather than to resolve taxonomic questions within *Saxicola*. Further work should expand and balance the sampling of species, individuals and geographic regions, with improved documentation of individual identity and recording provenance wherever possible. Integrating acoustic measurements with genetic and morphological data would then allow the observed patterns to be evaluated against independently derived evidence of population differentiation and evolutionary relationships. This would provide a basis for assessing the contribution of quantitative acoustic characterisation to the study of species boundaries and diversification within the genus.

## 6 CONCLUSIONS

This study demonstrates the feasibility of phase-based frequency tracking for the quantitative characterisation of alarm calls in the genus *Saxicola*. The extracted acoustic features reveal structured patterns of variation among calls, providing a basis for comparative analysis using principal component analysis and robust principal component analysis. Although the observed acoustic relationships offer opportunities to investigate species differentiation, geographic variation and hypotheses concerning evolutionary relationships, their biological interpretation is constrained by uneven sampling and uncertainty in individual and taxonomic identification. The method therefore provides a foundation for further comparative acoustic research, with integration of more comprehensive acoustic sampling and independent genetic evidence offering a promising direction for future investigation.

## 7 DATA AVAILABILITY

The recordings analysed in this study are part of the publicly available Xeno-Canto dataset.

## APPENDIX A: PHASE-BASED FREQUENCY TRACKING

### A.1 Phase-based frequency tracking: Derivation

Down-conversion to baseband, also known as ‘heterodyning’, is the process of translating a signal centred around a reference frequency to a frequency centred near 0 Hz, referred to as the *baseband*. This representation allows the subsequent signal processing to operate on the residual amplitude and phase evolution relative to a selected reference frequency.

Given a real bird call recording *x*(*t*), mixing with a reference carrier shifts the frame of reference into a zero-centred *baseband* domain. In this context, the baseband representation expresses the instantaneous frequency as a deviation relative to a static reference carrier, *f*_anchor_. The down-conversion yields the raw complex baseband components before filtering:

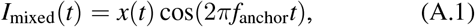

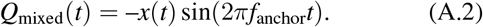

For a sinusoidal component

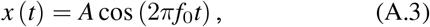

mixing with the anchor frequency gives

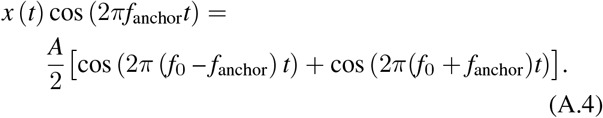

The low-pass filter removes the sum-frequency component, leaving the in-phase and quadrature baseband components, whose magnitude is therefore *A*/2. The instantaneous amplitude is consequently recovered as

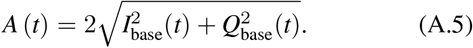

Following low-pass FIR filtering and window selection, we obtain the localized discrete vectors *I*_*k*_ and *Q*_*k*_, while maintaining the original sampling rate *f*_*s*_. The low-pass filtered in-phase *I*_*k*_ and quadrature *Q*_*k*_ components form a complex baseband signal,

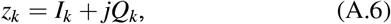

which may also be expressed in polar form as

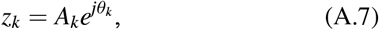

where *A*_*k*_ is the instantaneous amplitude and *θ*_*k*_ is the base-band phase.

The phase increment between successive samples can be obtained by considering the product of the current complex sample and the complex conjugate of the previous sample, 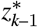. In exponential form,

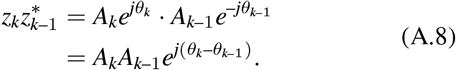

The argument of this complex product therefore corresponds directly to the phase increment between *z*_*k*–1_ and *z*_*k*_:

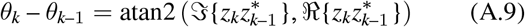

Using Eq. A.6, the real and imaginary components of the complex product can be written as

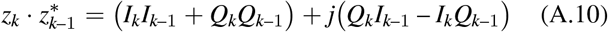

The resulting phase increments can then be converted to instantaneous frequency *f*_inst_ at sample *k* according to

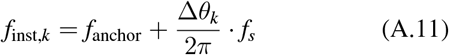

where *f*_anchor_ is the reference carrier frequency used for digital down-conversion and *f*_*s*_ is the sampling frequency.

The unwrapped baseband phase can subsequently be reconstructed by cumulative summation of the successive phase increments:

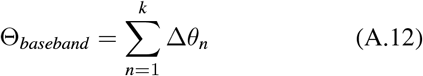

Because the phase increment is calculated between successive samples, explicit unwrapping of the cumulative phase is not required. The relevant constraint is instead that the phase change between adjacent samples remains within the principal interval (–*π, π*). Down-conversion reduces the phase increment from that associated with the absolute instantaneous frequency to that associated with its deviation from *f*_anchor_, thereby increasing the margin relative to this ambiguity limit. The estimator thus operates on:

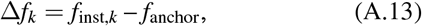

so the relevant baseband phase increment is

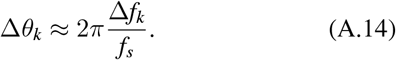

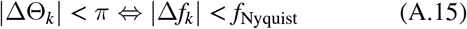

An important benefit of down-conversion is thus that the anchor frequency creates a large phase-increment safety margin around the frequency trajectory of interest.

### A.2 Synthetic tests

The performance of the phase-based frequency tracking algorithm was evaluated using two types of synthetic signals. The first consisted of a tapered linear chirp with an instantaneous frequency increasing from 4 kHz to 8 kHz. The second consisted of a tapered constant-frequency signal to which a 400 Hz frequency step of 1 ms duration was added, resulting in a box-shaped frequency excursion. The chirp evaluates the ability to follow a continuously changing frequency, while the box-shaped frequency jump evaluates temporal localisation and the ability to resolve a rapid frequency change. Both signals were evaluated under noise-free conditions and with random noise added. The noise-free synthetic chirp result is shown in Figure A.1. Both the Short-Time Fourier Transform (STFT) estimate and the phase-based frequency-tracking estimate are in close agreement with the theoretical instantaneous-frequency trajectory. This demonstrates that both methods can accurately recover the continuously varying frequency of the synthetic chirp under ideal, noise-free conditions.

The STFT was calculated using a window length of 36 samples at a sampling rate of 44 kHz, with an overlap of 75%. Increasing the window length produces a smoother frequency estimate due to increased averaging, but this comes at the cost of reduced temporal resolution. Figure A.2 shows the result for the same chirp with Gaussian random noise added with a standard deviation of 0.5. Both the STFT and phase-based estimates show the effect of the added noise; nevertheless, both continue to follow the theoretical frequency trajectory closely. The top panel compares the Hilbert envelope, calculated using (blue line), with the instantaneous amplitude calculated using the phase-based method. The Hilbert result comes out noisier: The default parameters for the band-pass filter included in the `scipy.signal.envelope` function are set wide, as no assumption can be made a priori on the frequency content of the signal. Figure A.3 shows the noise-free signal containing the box-shaped frequency excursion. The 400 Hz frequency jump with a duration of 1 ms corresponds to a time–bandwidth product of *BT* = 400 × 0.001 = 0.4. This value is close to the typical lower limit associated with a Gaussian pulse, making this a useful test case for evaluating the temporal and frequency resolution of the two methods. In contrast to the continuously varying chirp, this signal evaluates the ability of the frequency-tracking methods to resolve a short-duration, abrupt change in frequency. Figure A.4 shows the corresponding results with Gaussian random noise added with a standard deviation of 0.25.

### A.3 Discussion

Testing the STFT and phase-based methods using a box function with a time-bandwidth product of (*BT* = 0.4) shows comparable resolution performance. For the STFT, improved temporal resolution requires a shorter analysis window, which reduces frequency averaging and consequently increases sensitivity to noise. For the phase-based method, noise suppression is controlled through the Kalman filtering of the instantaneous-frequency trajectory; stronger smoothing improves noise robustness at the expense of temporal resolution. Thus, both approaches exhibit an inherent trade-off between noise suppression and temporal resolution, although this trade-off is controlled through different parameters.

In the phase-based approach, temporal resolution is not intrinsically determined by an analysis window. The raw instantaneous-frequency estimate is obtained at the sampling rate, while the effective resolution is subsequently controlled by the degree of Kalman filtering.

This means that the phase-based method separates the measurement from the resolution-control stage, whereas the STFT couples the two through the window function.

The choice of (*f*_anchor_) provides an additional degree of control that is not intrinsic to the STFT window. By selecting (*f*_anchor_) within the frequency band of the target signal, the target component is shifted towards baseband, while signals from other species occupying different frequency bands remain offset from zero frequency. The subsequent low-pass filtering can therefore attenuate these out-of-band components before the phase-based instantaneous-frequency estimate is calculated. This provides a means of reducing interference from other species while retaining the temporal resolution of the phase-based estimator.

For most signals, a fixed (*f*_anchor_) is sufficient. For signals exhibiting large frequency excursions, an optional second iteration employs the smoothed instantaneous-frequency estimate as a time-varying anchor. Although initially introduced to accommodate strongly frequency-modulated signals, this adaptive formulation may also provide a means of characterising call-to-call and intra-species frequency variation.

### A.4 Phase-based frequency tracking: Implementation

The signal processing pipeline executes in five sequential mathematical layers:

1. **Baseline Estimation:** A reference frequency *f*_anchor_ is estimated by counting zero-crossings in a short window around the maximum energy of the transient.
2. **Digital Down-Conversion (DDC):** The raw, real-valued input sequence *x*[*n*] is mixed with a complex local oscillator set explicitly to *f*_lo_ = *f*_anchor_, shifting the signal of interest down to a 0 Hz baseband center.
3. **Low-Pass Filtering & Local Window Extraction:** A sharp Finite Impulse Response (FIR) filter isolates the drift band, and an adaptive energy detector extracts the transient call boundaries without altering data dimensionality.
4. **Complex-Conjugate Phase Tracking:** Sample-to-sample phase variations are calculated via complex-conjugate cross-multiplications, bypassing unstable derivatives or hard-thresholded phase unwrapping functions.
5. **Kalman filter:** The instantaneous-frequency trajectory was smoothed using a forward–backward Kalman filter. In contrast to a causal forward Kalman filter, the forward– backward formulation uses information from both temporal directions. The bidirectional filtering constrains slowly varying frequency deviations using information from the complete transient, thereby reducing apparent drift while retaining control over the temporal smoothing through the Kalman process-noise parameter.

### A.5 Summary

Digital down-conversion was used to translate the frequency band of interest towards baseband prior to phase-based frequency tracking. The principal advantage is that the goniometric phase-difference estimator operates on the residual phase evolution relative to an explicit anchor frequency, (*f*_anchor_), rather than on the full carrier phase. Consequently, the requirement that the phase increment between successive samples remains within (–*π, π*) is more readily satisfied, providing increased margin against phase ambiguity and noise-induced branch crossings.

The choice of (*f*_anchor_) also provides explicit control over the frequency band being tracked. Components occupying substantially different frequency bands, such as signals from other species, remain offset from baseband and can therefore be attenuated by the subsequent low-pass FIR filter. The anchor frequency can consequently be selected according to a species-specific frequency range or estimated from a user-defined portion of the signal.

Rather than explicitly calculating and unwrapping the phase at each sample before differentiating it, the phase increment between consecutive complex samples is obtained directly from their relative angle. These sample-to-sample phase increments provide the instantaneous-frequency trajectory at the sampling rate. A forward–backward Kalman filter is then used to suppress noise while retaining the temporal evolution of the frequency trajectory. The degree of filtering determines the trade-off between noise suppression and temporal resolution, while the bidirectional formulation reduces directional and initialization effects and was found to minimise apparent frequency drift.

For signals exhibiting large frequency excursions, an optional second iteration can be performed in which the smoothed frequency trajectory obtained from the first pass is used as a time-dependent anchor frequency. This adaptive down-conversion keeps the residual frequency closer to baseband and allows strongly frequency-modulated signals to remain within the effective FIR bandwidth. In addition to extending the usable frequency range of the method, the resulting frequency trajectories provide a representation in which within-call dynamics and call-to-call frequency variation can be characterised.

## APPENDIX B: SEGMENTATION AND FEATURE-VECTOR CONSTRUCTION

### B.1 Introduction

Short transients, like bird calls present a particular challenge for time–frequency analysis. Their duration may be comparable to the time scale over which their spectral content changes, such that a method must simultaneously resolve rapid temporal variations and distinguish closely spaced frequency components. The short duration of these signals also limits the amount of temporal averaging that can be used to suppress measurement noise.

A conventional approach is the short-time Fourier transform (STFT), in which the length of the analysis window determines the temporal and spectral resolution. A shorter window improves temporal localization but provides less frequency averaging and is therefore more sensitive to noise. In contrast, a longer window improves frequency resolution and noise robustness at the expense of temporal resolution. Thus, the choice of window represents an inherent trade-off between temporal resolution and noise suppression.

An alternative is to estimate the instantaneous frequency directly from the phase evolution of the signal. Rather than determining the frequency content within a finite analysis window, the phase-based approach considers the phase increment between successive samples. For a sufficiently sampled oscillatory signal, this phase increment provides a direct estimate of the instantaneous frequency. The resulting estimate is initially obtained at the sampling rate, after which its temporal smoothness and noise sensitivity can be controlled independently through filtering.

The objective of the proposed phase-based frequency tracking method is not to assign a single characteristic frequency to a bird call, but to recover its time-dependent frequency trajectory. This distinction is important because species-specific information may be contained not only in the absolute frequency of a call, but also in how its frequency evolves throughout the transient. The method therefore treats frequency as a dynamic quantity (f(t)), allowing both within-call frequency modulation and variation between calls to be characterised.

In the present approach, the phase-based frequency estimation is preceded by digital down-conversion (DDC), in which the signal is mixed with a reference frequency, here denoted (*f*_anchor_). This shifts the frequency band of interest towards base-band, such that the phase increment represents the frequency deviation relative to the selected anchor frequency. The anchor therefore provides an explicit frequency reference that can be selected according to the expected frequency band of the signal. In recordings containing signals from multiple species, this also provides a means of reducing interference from signals occupying substantially different frequency bands: components distant from (*f*_anchor_) can be attenuated by the subsequent low-pass filtering.

The instantaneous-frequency trajectory obtained after down-conversion is subsequently smoothed using a forward– backward Kalman filter. The bidirectional formulation makes use of information from the complete transient rather than restricting the estimate at each sample to preceding observations. This reduces directional and initialization effects and, in practice, reduces apparent drift in the estimated frequency trajectories. The Kalman process-noise parameter provides control over the degree of smoothing, establishing a noise-suppression versus temporal-resolution trade-off analogous to that imposed by the STFT window, but without defining the instantaneous-frequency estimate through a fixed analysis window.

For signals exhibiting large frequency excursions, the fixed-anchor formulation can approach the limits imposed by the baseband filter bandwidth. An optional second iteration therefore uses the smoothed frequency trajectory from the first pass as a time-dependent anchor. The down-conversion then follows the estimated frequency trajectory, keeping the residual frequency closer to baseband and allowing strongly frequency-modulated signals to be analysed without requiring an excessively wide low-pass filter. Although this adaptive down-conversion was initially introduced to accommodate strongly varying synthetic signals, it also provides a potential means of characterising frequency variation between calls and within a species.

The resulting method therefore separates several aspects of the estimation problem that are coupled in conventional windowed time–frequency analysis: the frequency reference is controlled through (*f*_anchor_), frequency isolation through the baseband filter, and temporal smoothing through the Kalman filter. This provides a flexible framework for estimating and comparing the frequency trajectories of short bird call transients while retaining sensitivity to both rapid within-call dynamics and longer-term variation between calls.

### B.2 Feature vector construction

Each instantaneous-frequency trajectory was characterized by dividing the transient into onset, sustain, and decay components. The onset was defined from the first crossing of a threshold corresponding to a fixed fraction of the maximum envelope amplitude. The end of the transient was defined by the first subsequent crossing of a noise-floor threshold, estimated from the terminal portion of the envelope. The sustain was defined as a constant-amplitude region surrounding the maximum amplitude, rather than as a frequency-defined region. Its lower amplitude boundary was varied systematically, and the segmentation giving the minimum global standard error of the fitted frequency trajectory was selected. Thus, the segmentation was determined jointly from the amplitude envelope and the frequency trajectory.

Within the resulting segments, frequency was modelled as a function of normalized amplitude. The onset and decay portions were fitted using polynomials, whereas the sustain frequency was represented by its mean value. For each segment, the fitted frequency models were subsequently used to derive descriptors of the frequency trajectory. These included the normalized frequency excursions, signed slopes, and normalized curvatures of the onset and decay, together with the maximum frequency, relative sustain frequency, and frequency variability within the sustain. The slope was evaluated at the midpoint of the corresponding amplitude interval, while curvature was obtained from the second derivative of the fitted polynomial and normalized by the squared amplitude range and the magnitude of the corresponding frequency excursion:

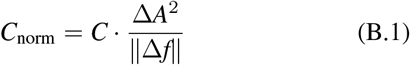

**FIGURE B.1.**
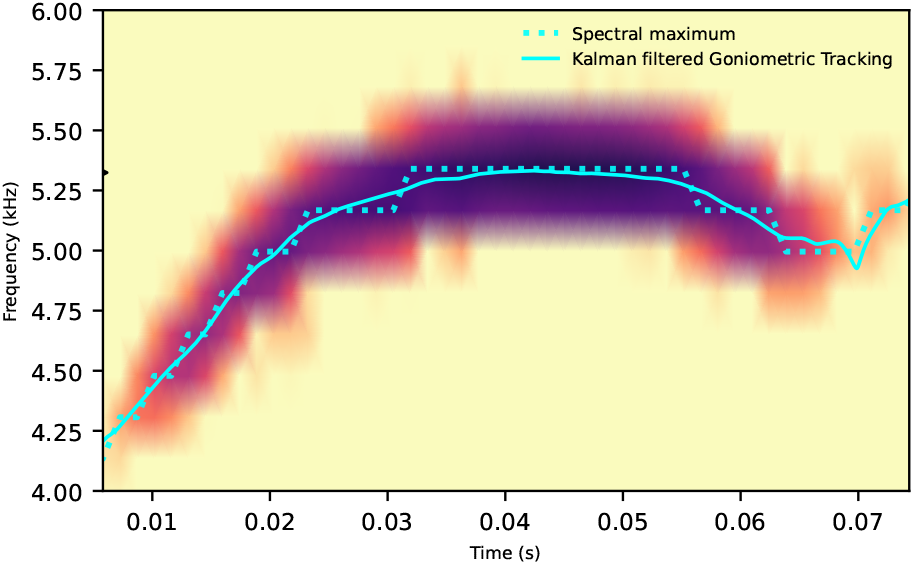
Sparse modal representation of a synthetic transient repeated three times to form a click sequence with a 3 ms interclick interval. The transient is made by adding three damped resonances (frequency/quality factor: 40 kHz/6, 100 kHz/10 and 190 kHz/20). The dictionary is built from three estimated damped resonances, using, from left to right, 4, 14, or 24 atoms.

Temporal morphology was characterized by the total transient duration and the duration for which the envelope exceeded 50% of its maximum amplitude. The resulting descriptors form a feature vector representing the frequency, amplitude–frequency geometry, and temporal structure of each transient. Although residual errors from the segment fits were calculated, these were retained as diagnostic measures and were not included in the final clustering feature vector.

Each bird call was represented by its instantaneous frequency and amplitude envelope as a function of time. The transient was subsequently divided into three regions—onset, sustain, and decay—to describe its temporal and spectral morphology. The segmentation was driven primarily by the amplitude envelope, while the frequency trajectory was used to determine the optimal boundaries between the three regions.

The onset was defined from the first crossing of a threshold corresponding to 10% of the maximum envelope amplitude. Following the maximum amplitude, the end of the transient was defined by the first crossing below an estimated noise level. The noise level was calculated from the final portion of the signal as the mean amplitude plus twice its standard deviation. The sustain region was then determined by systematically varying its lower amplitude boundary. For each candidate boundary, the onset and decay frequency trajectories were fitted and the sustain frequency was represented by its mean value. The segmentation yielding the minimum global standard error of the frequency fits was retained.

A key aspect of this procedure is that the onset and decay trajectories were characterized as frequency as a function of amplitude, rather than frequency as a function of time. Thus, for the onset and decay regions, the instantaneous frequency was represented as

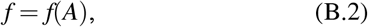

where *A* is the instantaneous amplitude envelope. Polynomial functions were fitted separately to the onset and decay regions. This representation describes the geometry of the frequency trajectory relative to the amplitude evolution of the transient, rather than its temporal evolution alone. It also provides a common amplitude-based coordinate system for comparing transients with different durations or rates of amplitude change.

From these fitted trajectories, descriptors of frequency excursion, slope, and curvature were calculated. The slope was evaluated at the midpoint of the corresponding amplitude interval, providing a measure of the local frequency change with amplitude. For a quadratic fit,

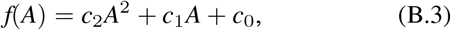

the corresponding slope and curvature are

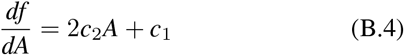

and

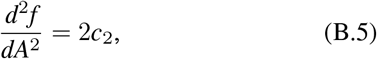

respectively. Frequency excursions and slopes were normalised by the maximum frequency, while curvature was scaled by the squared amplitude range and the magnitude of the associated frequency excursion. This normalization reduces the dependence of the descriptors on absolute frequency and amplitude scale while retaining the shape of the frequency– amplitude trajectory.

The final feature vector comprised ten descriptors:

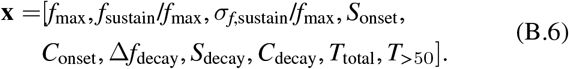

Here, *f*_max_ is the maximum instantaneous frequency; the second and third terms describe the mean and variability of frequency during the sustain region; *S*_onset_ and *S*_decay_ describe the normalized slopes of the onset and decay trajectories; *C*_onset_ and *C*_decay_ describe their normalized curvature; and Δ*f*_decay_ describes the normalized frequency excursion over the decay. The final two features represent the total transient duration and the duration for which the envelope exceeded 50

Several initially derived descriptors were excluded from the final feature vector to obtain a more compact representation. These included the onset frequency excursion, the sustain amplitude range, and the relative durations of the onset, sustain, and decay regions. The remaining features were standardized before principal component analysis (PCA).

## APPENDIX C: XENO CANTO

**TABLE C.1.**
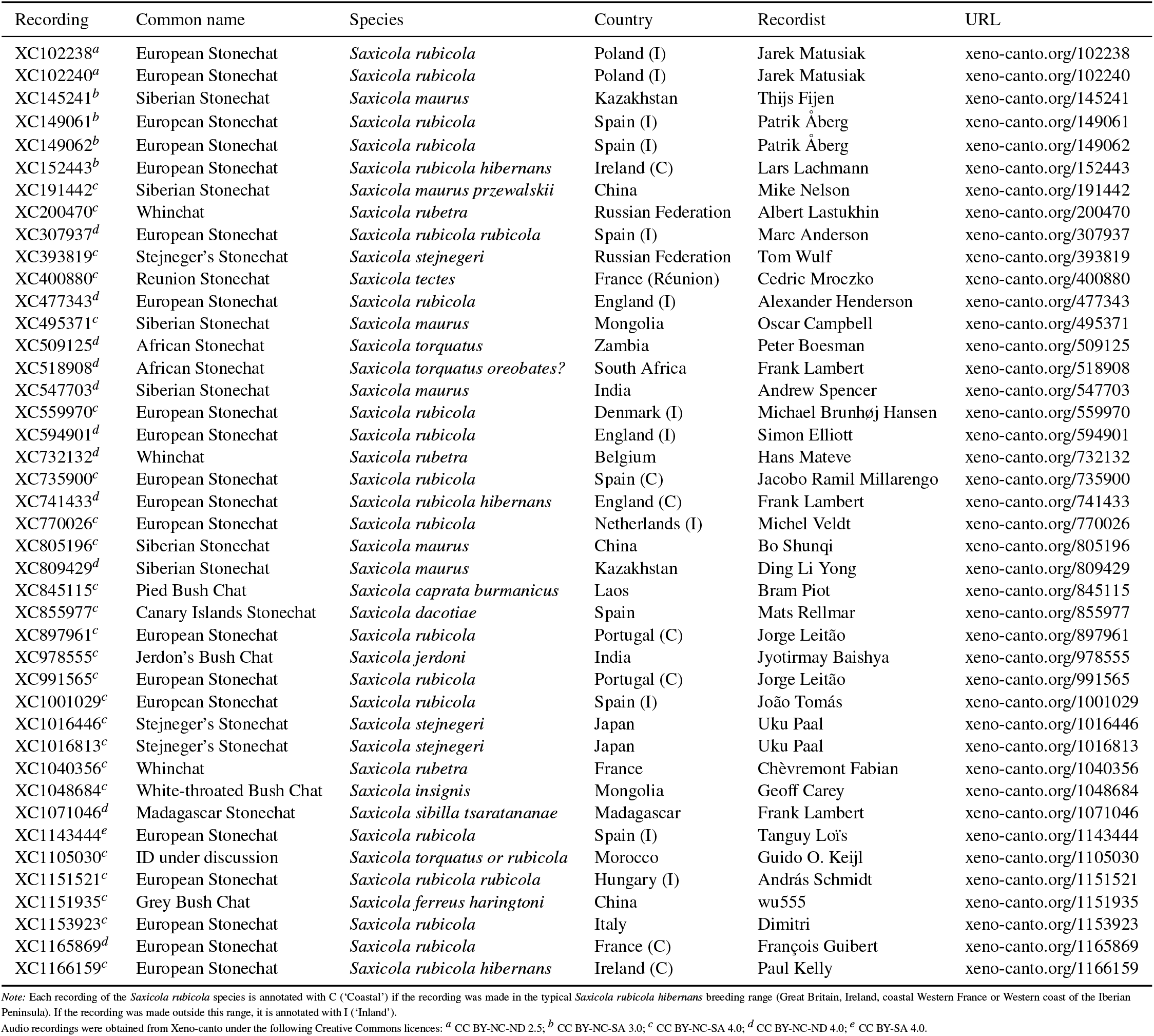
Recordings included in the *Saxicola* dataset.

